# Chemical and Sensory Analyses of Cultivated Pork Fat Tissue as a Flavor Enhancer for Meat Alternatives

**DOI:** 10.1101/2024.05.31.596657

**Authors:** Emily T. Lew, John S.K. Yuen, Kevin L. Zhang, Katherine Fuller, Scott C. Frost, David L. Kaplan

## Abstract

The emerging field of cellular agriculture has accelerated the development of cell-cultivated adipose tissue as an additive to enhance the flavor of alternative meat products. However, there has been limited research to evaluate the sensory profile of *in* vitro-grown tissues compared to conventionally obtained animal fat. This study aimed to investigate the aromatic characteristics of cell-cultivated fat tissue as a flavor enhancer for meat alternatives. Porcine dedifferentiated fat cells were clonally isolated and differentiated into adipocytes. This cultured adipose tissue was then analyzed alongside native porcine fat using gas chromatography-mass spectrometry (GC/MS) coupled with descriptive sensory analysis by human panelists. This evaluation enabled quantitative and qualitative assessments of volatile compounds released during cooking for both in vitro and in vivo porcine fats. The volatile profiles generated during the cooking process and fatty aroma characteristics reported by sensory panelists were largely similar between the two fat sources, with some differences in the concentration of select compounds and aroma attributes. Ultimately, the panelists found comparable overall liking scores reported between the conventional and cultured porcine fats. These findings provide valuable sensory evidence supporting the viability of cell-cultivated adipose tissue as a flavor component of meat alternatives, substituting for conventional animal fat.

## Introduction

The increasing global population and the environmental, ethical, and health-related concerns associated with traditional livestock farming have accelerated the search for sustainable and humane alternatives to industrialized meat production ^1–6^. Currently, plant-based meats made from soy, tofu, and textured vegetable proteins serve as alternative protein sources for vegetarians, vegans and other consumers interested in reducing their meat consumption. Similarly to cultivated meat, plant-based proteins have a much lower environmental impact compared to animal-derived proteins due to lower water use, land use and reduced greenhouse gas emissions ^7^. Although many consumers are interested in plant-based meats, taste is a primary barrier to consumption due to flavor differences compared to conventional animal proteins ^8^. In response to these challenges, recent advances in cellular agriculture have demonstrated the potential for producing cell-cultured adipose tissue, or fat, as a key component in replicating the sensory qualities and nutritional value of meat products ^9–12^. However, while the theoretical advantages of cell-cultivated fat are widely discussed, limited empirical evidence exists and there remains a lack of comprehensive evaluations supporting the sensory characteristics and overall viability of cultivated fat as a meat alternative for traditional livestock-generated fat. ^13,14^

Despite the limited presence in the consumer marketplace, many studies have been conducted to examine the potential for consumer acceptance of cultivated meat ^15–17^. In these studies, sensory characteristics similar to conventional meat is one of the primary determinants of consumer acceptance ^18,19^. Additionally, several reviews have been published which outline the hypothetical acceptance of cultivated meat, but do not discuss measurable sensory qualities of cultivated meat aroma or flavor ^13,20–22^. A handful of sensory evaluation studies including real cultivated meat samples have been conducted which determined that the cultivated meat products closely resembled conventional analogues (Benjamison et al., Ong et al., Lee et al. and Patiska et al.). Although the findings of these studies demonstrate the potential for cultivated meat to emulate conventional animal products, issues include low numbers of human panelists and comparisons that were conducted to nonequivalent conventional products ^13^. To address these gaps, we present a study with over 50 participants using a comparable target conventional meat product to evaluate the sensorial qualities of cultivated porcine adipose.

In our previous research, adipocytes were grown at a laboratory scale and mechanically aggregated into macroscale tissues resembling native pig fat ^23^. These *in vitro*-grown porcine adipocyte tissues demonstrated similar fatty acid composition to native pig fat, suggesting the potential similarity in taste and aroma characteristics with native porcine adipose. Using lipidomic analysis, it was found that *in vitro*-grown and native porcine fats contained similar fatty acid profiles, especially intracellular triacylglyceride fractions. Significant differences were found in the phospholipid fraction where cultivated porcine fat contained lower amounts of 18:2 and 18:3 fatty acids and higher amount of 20:1. Media supplementation with Intralipid remedied this difference, whereby an increase of 18:2 and 18:3, and a decrease of 20:1 was found. Additionally, *in vitro*-grown and native porcine fats shared similar ratios of monounsaturated fatty acids, polyunsaturated fatty acids and saturated fatty acids with the Intralipid supplementation.

Building upon these previous findings, the goal of the present study was to evaluate the composition and sensory characteristics of cell-cultivated porcine adipose tissue in detail, towards the evaluation of this process for flavor-enhancing components in alternative meat products. To achieve this objective, porcine fat was generated in vitro using a porcine dedifferentiated fat (PDFAT) cell subclone produced via single cell seeding from the original mixed population of primary DFAT cells. Gas chromatography-mass spectrometry (GC/MS) paired with human sensory evaluation were utilized to quantitatively and qualitatively assess volatile compounds released from both the *in vitro* and *in vivo* porcine fats during heating (cooked fat aroma). Additional GC/MS analyses using an olfactory detection port (ODP) was utilized to identify specific compounds responsible for various porcine fat characteristics reported during the human sensory trials.

## Results

### Study Design

**Figure 1** shows an overview of the study design including isolation of a clonal population of PDFAT, macroscale aggregation of cultivated fat and use of fat samples in sensory workflow (**Figure 1**).

**Figure 1.**
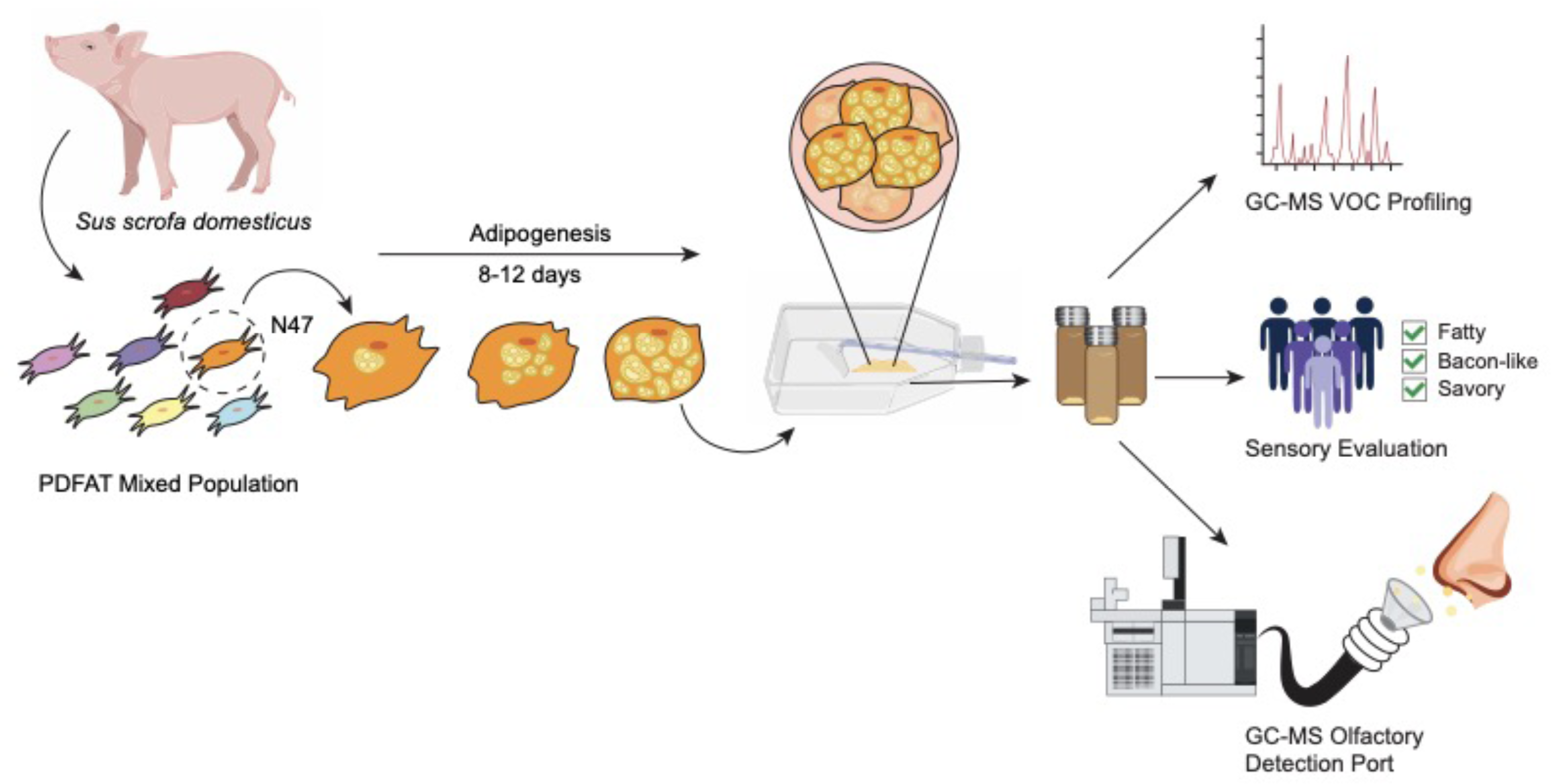
Graphical overview of the methodology. A clonal population of porcine dedifferentiated fat cells were isolated, mechanically aggregated via cell scraping and used for GC/MS and sensory analysis.

### Clonally isolated PDFAT cells optimized for lipid accumulation were aggregated into 3D fat samples

To obtain a pure cell line from previously isolated PDFAT, single cell sorting was used to separate single cells. The goal was to select a high preforming clone and optimize adipogenesis for a variety of differentiation media components (**Figure 2A**). A total of 576 single cells were seeded into regular culture media (N), 50% conditioned media (C) and vitronectin supplemented media (V) in 96-well plates. At 24 hours after seeding, wells were screened for successful single cell seeding indicated by isolated cell populations. Clonal populations were grown to confluence and visually screened for proliferation. Intracellular lipid accumulation of all viable clones was quantified with Oil Red O. Five clones N18, N20, N47, C15, and C34 were selected as they were found in the top 15 clones of the lipid accumulation screen (**Supplementary Fig. S1**) while previously exhibiting strong proliferation based on visual observation (data not shown). These five clones were monitored for six passages before reducing the screen to clones N18, N47, and C34). Clones N18, N47 and C34 were selected from the top five clones based on visual proliferation screening and lipid accumulation (data not shown). Clones N18, N47 and C34 were screened for proliferation through passage nine (**Supplementary Fig. S2**). Due to higher lipid accumulation, long-term culture continued with clone N47 (**Figure 2B, i**). Across 100 passages, clone N47 had an average doubling time of 60.6 hrs. with an average of 1.15 doublings per passage (**Figure 2B; ii, iii**). Media optimization was conducted for adipogenesis agonists dexamethasone, rosiglitazone, 3-isobutyl-1-methylxanthine (IBMX), and BMP-4 to better inform media formulations for the mixed population PDFAT and clone N47. Standard adipogenic culture media contained both dexamethasone and rosiglitazone in adipogenic induction and lipid accumulation media and 0.5mM IBMX. Culture media was altered to omit rosiglitazone in lipid accumulation media and reduce IBMX concentration to 0.1mM thereafter (**Figure 2C; i, ii**). While a high degree of lipid accumulation was observed, lipid morphology remained multilocular with these optimizations (**Supplementary Fig. S3**). Additionally, the presence of the differentiation factor BMP-4 was tested at 20 ng/mL but did not yield higher lipid accumulation (**Figure 2C, iii**). Cells became lipid-laden within eight days of adipogenesis (**Figure 2D**). Approximately 1g of adipose tissue was harvested via cell scraping for subsequent downstream experiments (**Figure 2E**). With an initial seeding density of 11,000 cells/cm^2^, T175 flasks became confluent within approximately 4 days. One culture flask with 175 cm^2^ surface area yielded approximately 150 mg of cell mass eight days of adipogenesis. After harvest, scraped adipocytes were stored at −80°C for GC/MS and sensory analysis.

**Figure 2.**
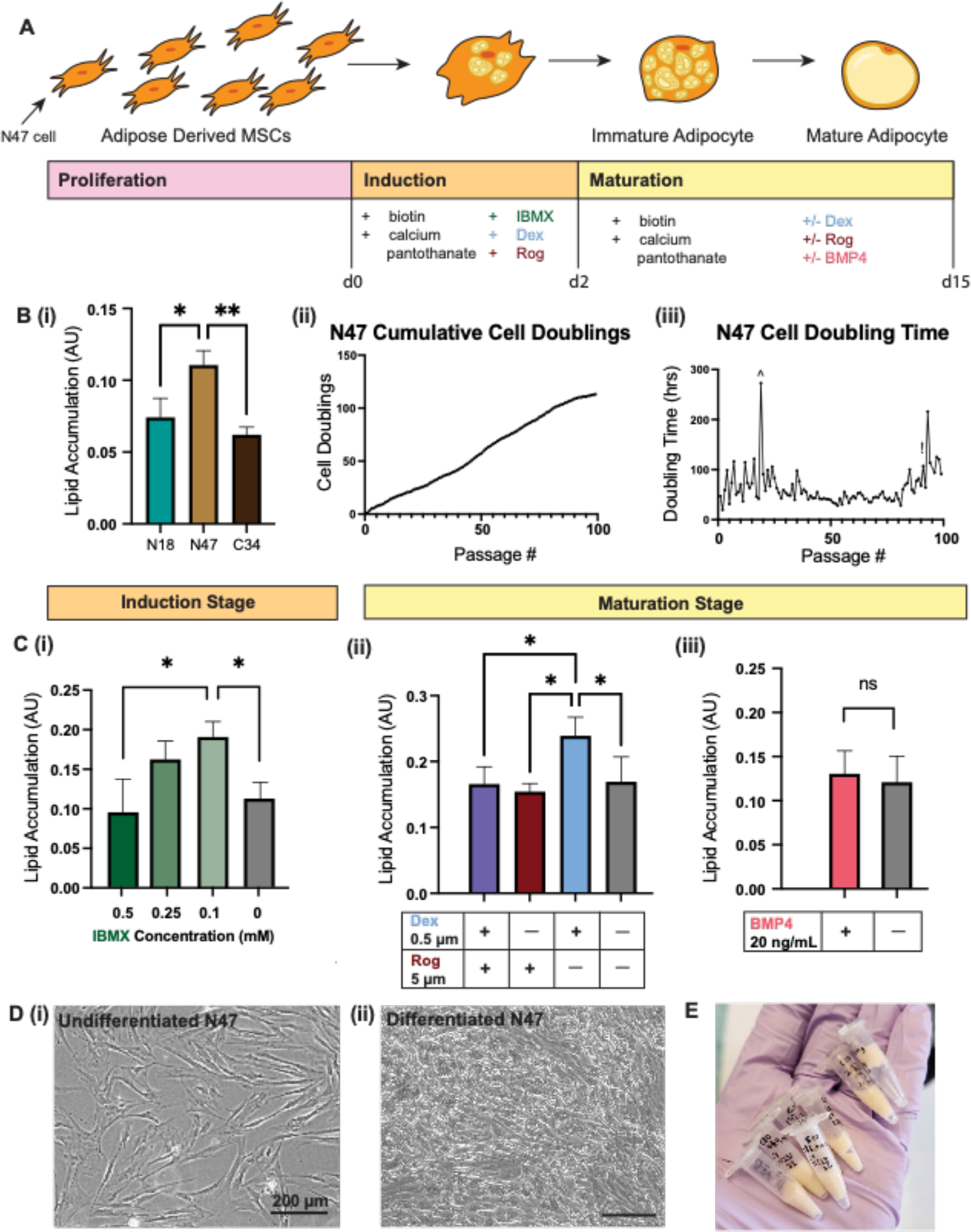
Single-cell PDFAT clones grown and optimized for cultured fat. (**A**) Schematic depicting stages of adipogenesis and media components targeted for optimization. (**B**) Three porcine DFAT clones (P12) screened for lipid accumulation after eight days of adipogenesis via Oil Red O staining. (**B, i**) Screening of clones N18, N47 and C34. (**B, ii)** Cumulative cell doublings. Data was analyzed with one-way ANOVA, where p≤ 0.05 (*) and p≤ 0.01 (**) (**B, iii**) Hours per doubling. (^) indicates passage with low cell count due to poor detachment and (!) indicates passage 95 reporting no growth (data point not shown). (**C**) Media optimization for adipogenic induction and maturation stages. N47 was screened for lipid accumulation after eight days of adipogenesis via Oil Red O staining. (**C, i)** Presence of IBMX in the first 2-3 days of adipogenesis (P12, n=3). Data was analyzed with one-way ANOVA, where p≤ 0.05 (*). (**C, ii**) Presence of dexamethasone and rosiglitazone in lipid accumulation media (P3). Both dexamethasone and rosiglitazone were present for the first 2-3 days of adipogenesis (induction media). Data was analyzed with two-way ANOVA, where p≤ 0.05 (*). (**C, iii**) Presence of BMP4 at 20 ng/mL in the lipid accumulation stage (n=3). Data was analyzed with unpaired t-test, where p≤ 0.05 (*). (**D**) N47 morphology before and after differentiation. (**D, i**) Undifferentiated and (**D, ii**) differentiated cells after eight days of adipogenesis. Scale bar 200 um. (**E**) N47 adipocytes were mechanically harvested with a cell scraper and aggregated into masses of cultured fat. Fat was aliquoted into portions of 150-200 mg in 0.6 ml Eppendorf tubes.

### Volatile chemistry of cooked porcine fat was analyzed using GC/MS detection

Comparison of total ion chromatograms of conventional and in vitro fat samples revealed very similar profiles, apart from additional low volatility (high retention) compounds found in conventional fat samples and additional high volatility (low retention) compounds from in vitro-grown fat samples (**Figure 3A**). Compounds identified by GC/MS in cultivated and conventional porcine fat were ranked by peak area. Comparison of the top 20 compounds present in cultivated and conventional porcine fat reveals that they share long and medium chain fatty acid compounds such as hexadecenoic acid, hexanoic acid, (E)-9-octadecanoic acid, octadecanoic acid, tetradecanoic acid, among others (**Figure 3B, Table 1, Table 2**). The complete list of 60 shared compounds identified in cultivated and conventional porcine fat can be found as Supplementary Table S1 online. Unique compounds identified in each fat sample can be found as Supplementary Table S2 and Supplementary Table S3 online.

**Figure 3.**
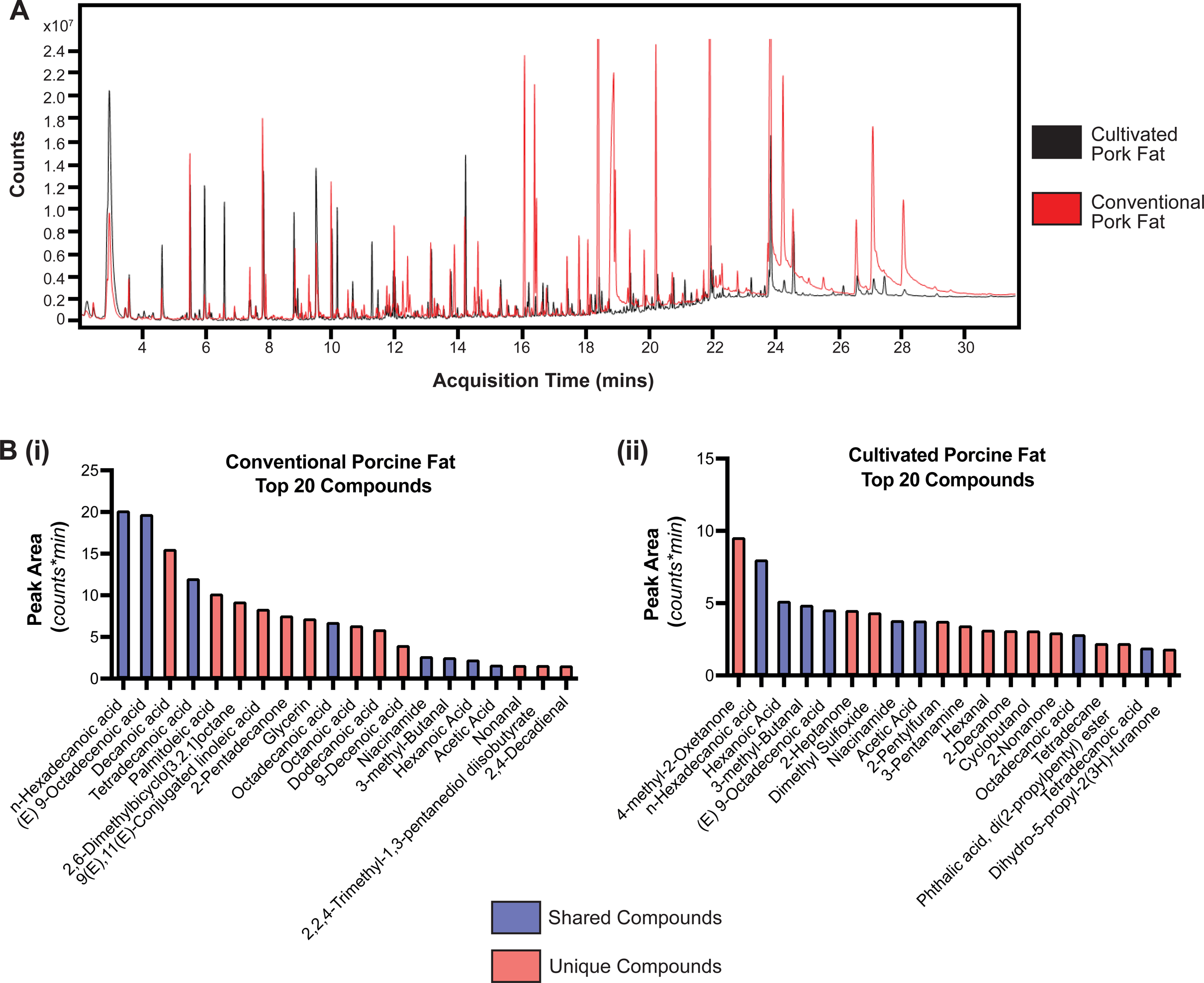
Volatile organic compound (VOC) profiling of conventional (livestock sourced) and cell cultivated porcine fat. (**A**) Total ion chromatograms of conventional pork belly fat (red) and N47 cultured porcine fat (black). Volatile and semi-volatile chemistry from each fat sampled using DHS with subsequent analysis using gas chromatography and mass spectrometry detection. Samples were heated to 120°C with 15 min incubation and 30 min trapping time. (**B**) Comparison of top 20 volatile compounds from (**B, i**) conventional pork fat and (**B, ii**) N47 cultivated porcine fat using peak area (n=3). Red denotes compounds unique to either conventional or cultivated fat samples. Blue denotes that are shared between conventional and cultivated fat samples.

**Table 1.** Top 20 volatiles by peak area from conventional porcine fat. “*” denotes compounds in the top 20 that were also detected with the GC/MS olfactory detection port. Tentative identification is indicated by “**” (n=3).

| Compound | Peak Area | Retention Time (min.) | RI (meas.) | RI (ref.) | Reference |
| --- | --- | --- | --- | --- | --- |
| n-Hexadecanoic acid | 20.138 | 23.900 | 2908 | 2910 | <sup>24</sup> |
| (E) 9-Octadecenoic acid | 19.700 | 27.148 | ** | 3173 | <sup>25</sup> |
| *Decanoic acid | 15.507 | 18.452 | 2272 | 2279 | <sup>26</sup> |
| Tetradecanoic acid | 11.990 | 21.986 | 2696 | 2712 | <sup>27</sup> |
| Palmitoleic acid | 10.155 | 24.297 | 2944 | 2960 | <sup>25</sup> |
| 2,6-Dimethylbicyclo[3.2.1]octane | 9.191 | 28.104 | ** | ** | ** |
| 9(E),11(E)-Conjugated linoleic acid | 8.319 | 28.113 | ** | 3168 | <sup>28</sup> |
| *2-Pentadecanone | 7.523 | 16.125 | 2027 | 2011 | <sup>29</sup> |
| Glycerin | 7.168 | 18.850 | 2316 | 2322 | <sup>30</sup> |
| Octadecanoic acid | 6.738 | 26.617 | NA | 3181 | <sup>31</sup> |
| Octanoic acid | 6.353 | 16.434 | 2058 | 2070 | <sup>32</sup> |
| Dodecanoic acid | 5.855 | 20.277 | 2483 | 2502 | 24 |
| 9-Decenoic acid | 3.970 | 18.982 | 2332 | 2348 | 33 |
| Niacinamide | 2.637 | 24.603 | 2971 | 2960 | 25 |
| *3-methyl-Butanal | 2.504 | 2.907 | 852 | 876 | 30 |
| *Hexanoic acid | 2.231 | 14.224 | 1844 | 1849 | 26 |
| *Acetic acid | 1.612 | 10.428 | 1518 | 1465 | 34 |
| *Nonanal | 1.572 | 8.849 | 1395 | 1390 | 35 |
| 2,2,4-Trimethyl-1,3-pentanediol diisobutyrate | 1.568 | 14.695 | 1887 | 1591 | 36 |
| 2,4-Decadienal | 1.526 | 13.913 | 1815 | 1762 | 34 |

**Table 2.**
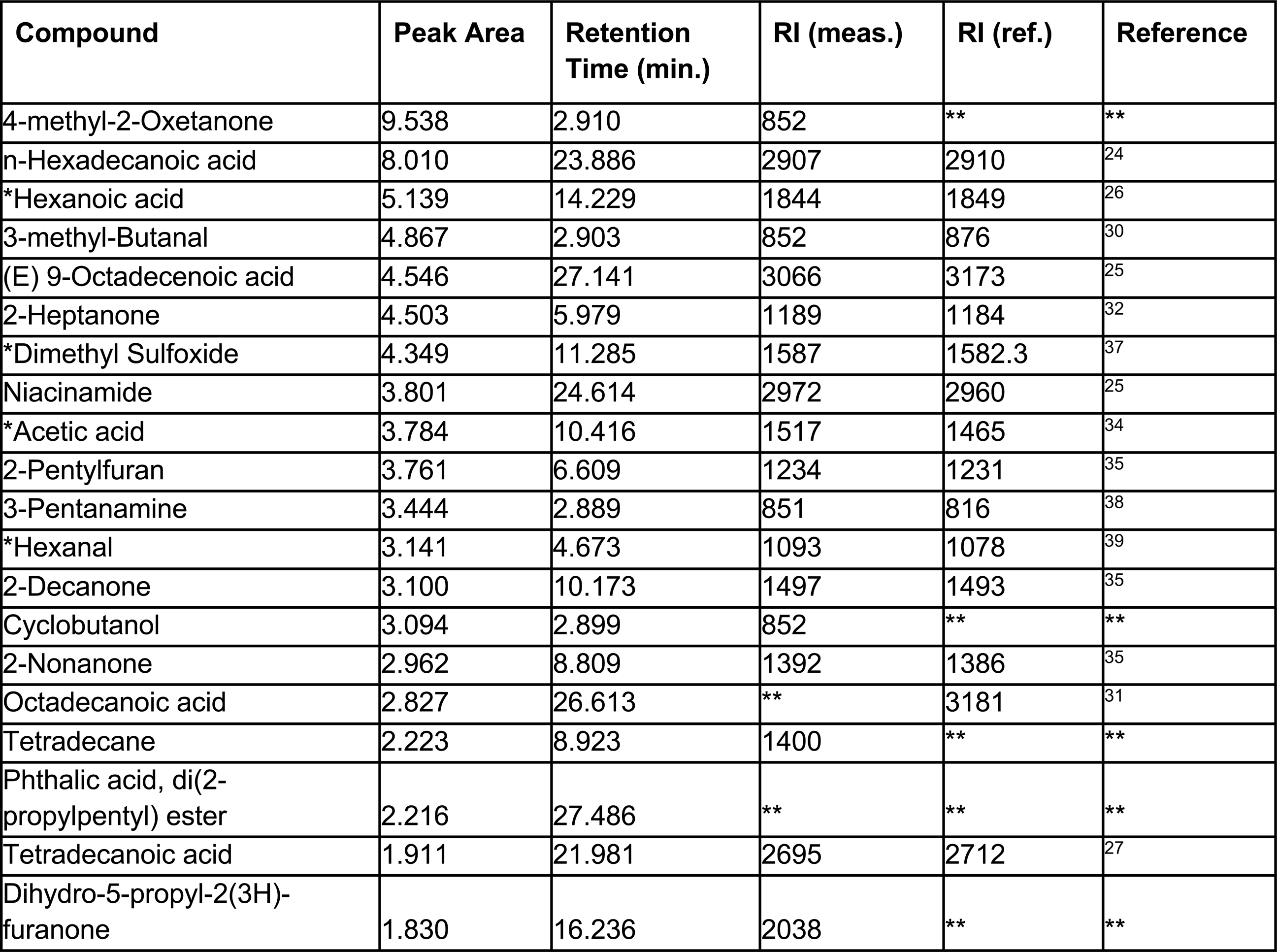
Top 20 volatiles by peak area from cultivated porcine fat. “*” denotes compounds in the top 20 that were also detected with the GC/MS olfactory detection port. Tentative identification is indicated by “**” (n=3).

| Compound | Peak Area | Retention Time (min.) | RI (meas.) | RI (ref.) | Reference |
| --- | --- | --- | --- | --- | --- |
| 4-methyl-2-Oxetanone | 9.538 | 2.910 | 852 | ** | ** |
| n-Hexadecanoic acid | 8.010 | 23.886 | 2907 | 2910 | 24 |
| *Hexanoic acid | 5.139 | 14.229 | 1844 | 1849 | 26 |
| 3-methyl-Butanal | 4.867 | 2.903 | 852 | 876 | 30 |
| (E) 9-Octadecenoic acid | 4.546 | 27.141 | 3066 | 3173 | 25 |
| 2-Heptanone | 4.503 | 5.979 | 1189 | 1184 | 32 |
| *Dimethyl Sulfoxide | 4.349 | 11.285 | 1587 | 1582.3 | 37 |
| Niacinamide | 3.801 | 24.614 | 2972 | 2960 | 25 |
| *Acetic acid | 3.784 | 10.416 | 1517 | 1465 | 34 |
| 2-Pentylfuran | 3.761 | 6.609 | 1234 | 1231 | 35 |
| 3-Pentanamine | 3.444 | 2.889 | 851 | 816 | 38 |
| *Hexanal | 3.141 | 4.673 | 1093 | 1078 | 39 |
| 2-Decanone | 3.100 | 10.173 | 1497 | 1493 | 35 |
| Cyclobutanol | 3.094 | 2.899 | 852 | ** | ** |
| 2-Nonanone | 2.962 | 8.809 | 1392 | 1386 | 35 |
| Octadecanoic acid | 2.827 | 26.613 | ** | 3181 | 31 |
| Tetradecane | 2.223 | 8.923 | 1400 | ** | ** |
| Phthalic acid, di(2-propylpentyl) ester | 2.216 | 27.486 | ** | ** | ** |
| Tetradecanoic acid | 1.911 | 21.981 | 2695 | 2712 | 27 |
| Dihydro-5-propyl-2(3H)-furanone | 1.830 | 16.236 | 2038 | ** | ** |

### Sensory evaluation panelists were able to discriminate between conventional and cultivated fat samples

Significant discrimination was shown between the two fat samples. 41 out of 54 participants were able to correctly discriminate between conventional (livestock sourced) and cultivated fat tissue. Panelists were presented with three fat samples in unique combinations containing either two cultivated and one conventional sample or one cultivated and two conventional samples. Despite the different sample presentations, enough panelists were able to correctly discriminate between the cultivated and conventional samples to demonstrate statistical significance. The critical number of correct responses required to demonstrate statistical significance (p<0.05) in a triangle test is 25/54 ^40^ (**Supplementary Table S4)**. The full summary of triangle test responses can be found in Supplementary Table S5.

### Meat-like aroma attributes identified by sensory evaluation panelists were largely similar for conventional and cultivated fats, while certain attributes were slightly more prevalent in cultivated fat samples

Panelists indicated their sensory perceptions of *in vivo* and *in vitro*-grown pig fat through a CATA task. An attribute list was generated using 18 common descriptors used in previously published sensory evaluations of meat products and aroma/flavor wheels ^41–44^. Overall panelist usage of the 18 attributes is shown as a stacked bar plot (**Figure 4A**). The total height of each bar corresponds to the total usage percentage of each descriptor. For example, the term “savory’’ was used 50% of the time by all panelists when evaluating cultivated pork fat. Between the two fat samples, 55 panelists were given the opportunity to check “fatty” a total of 110 instances, of which “fatty” was checked 73 times (66.4%). Additionally, each bar is labeled with the usage percentage of the attribute for a given fat sample calculated by the percentage of the 55 panelists. For example, 65.5% of panelists checked “fatty” for cultivated pig fat. For both fat samples, the attribute “fatty” was checked most of the time followed by “savory’’, “fried”, and “meaty” which were used 47.3%, 47.3%, and 38.2% of the time for cultivated fat samples. Term usage differed more greatly with negative attributes such as “musty”, “barnyard”, “cardboard” and “rancid’’, where panelists checked these attributes more often when evaluating cultivated fat samples. Panelists also check the “fried’’ attribute more often for the cultivated samples than the conventional samples.

**Figure 4.**
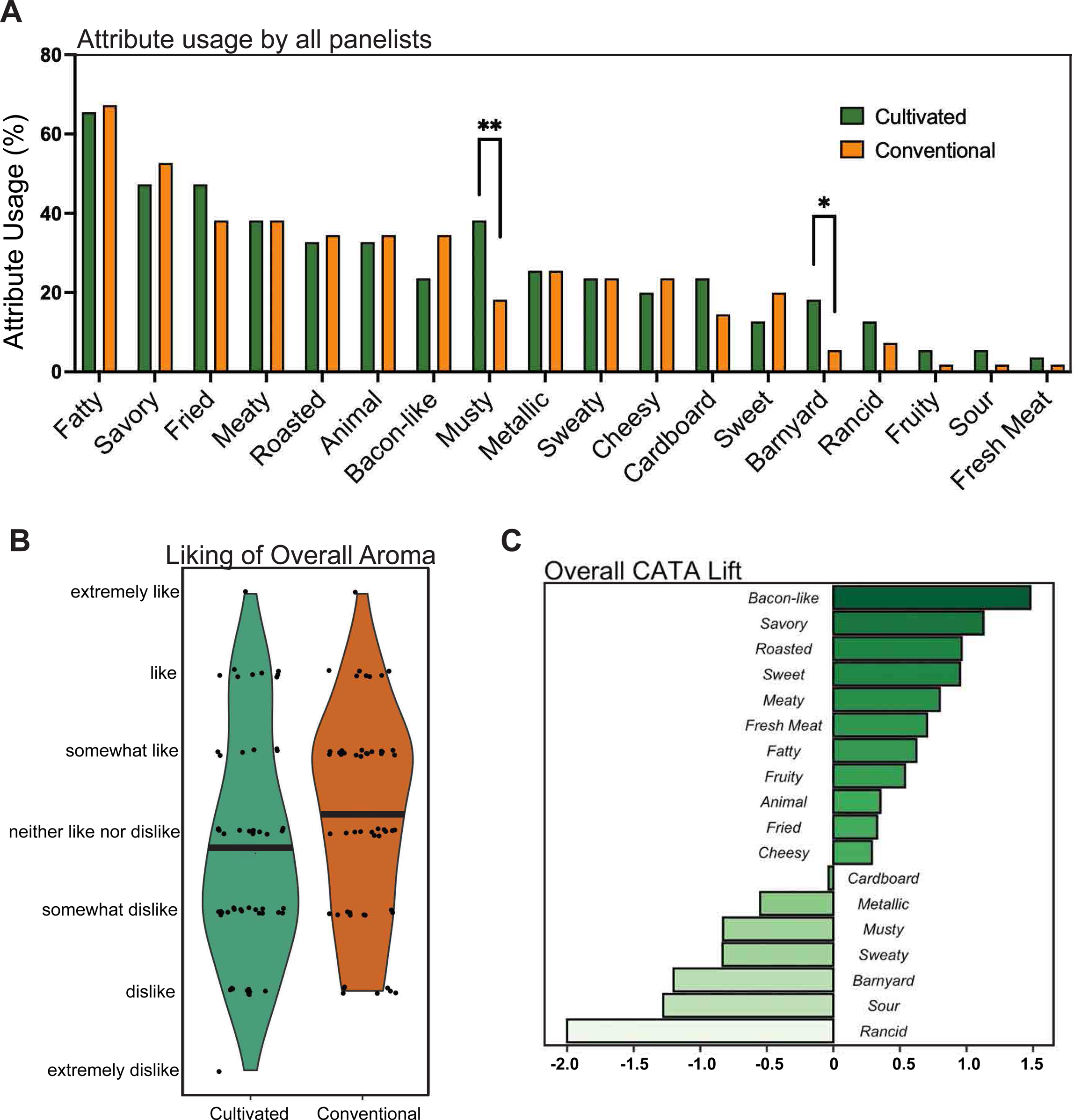
Aroma sensory evaluation conducted with 55 panelists. (**A**) Overall attribute usage by all judges in CATA study for cultivated and conventional porcine fat samples. Sample groups were compared with Fischer’s exact test, where p≤ 0.05 (*) and p≤ 0.01 (**). (**B**) Violin plot demonstrating distribution of overall liking of aroma between cultivated and conventional porcine fat samples on a 7-point hedonic scale of Extremely Dislike to Extremely Like. Mean values for each fat sample are shown by a solid black line. (**C**) Penalty/lift analysis relating degree of liking to specific sensory attributes in cultivated and conventional porcine fat samples.

To investigate the overall opinion of the aromas of the fat samples, panelists ranked the overall aroma of the samples on a 7-point hedonic scale from “Extremely Dislike” to “Extremely Like” (**Figure 4B**). The violin plot displays hedonic ratings for each fat sample. On average, panelists expressed no difference with respect to the aroma of both conventional and cultivated pig fat.

CATA-lift analysis was used to investigate the changes in the hedonic ratings when a specific attribute was detected compared with not detected (**Figure 4C**)^45^. Penalty/lift analysis shows the mean hedonic difference for each of the 18 attributes. For example, when the attribute “bacon-like” was checked for either cultured or conventional fat samples, a 1.5-point mean increase in overall liking was found. Liking increased when panelists detected “bacon-like”, “savory”, “roasted”, “sweet”, “meaty”, “fatty”, “fruity”, “animal”, “fried” and “cheesy” aromas, while liking decreased when panelists detected “cardboard”, “metallic”, “musty”, “sweaty”, “barnyard”, “sour” and “rancid” aromas.

### Major contributors to “musty” and “barnyard” aromas in both conventional and in vitro-grown fat samples include medium-chain saturated fatty acids

Out of 144 compounds detected in conventional pork fat, 32 were selected as aroma-active (**Table 3**). Sensory impactful compounds were of similar classification with the addition of ketones and benzene derivatives. Out of 179 compounds detected in cultivated pork fat, 26 compounds were selected as aroma-active (**Table 4**). These compounds include aldehydes, alcohols, acids, sulfur-containing compounds, and furans.

**Table 3.** GC/MS Olfactory Detection Port analysis of conventional pork fat. Compounds were tentatively identified using NIST database matching; compound confirmation was made using published retention index values. Tentative identification is indicated by “**” (n=3).

| Compound | Aroma | RI (meas.) | RI (ref.) | Reference |
| --- | --- | --- | --- | --- |
| 3-methyl-butanal | butter | 940 | 876 | 30 |
| Pentanal | butter, sweet | 975 | 979 | 32 |
| Hexanal | grass | 1048 | 1078 | 39 |
| 12,15-Octadecadiynoic acid, methyl ester | sour | 1095 | 2590 | 53 |
| Heptanal | sweet | 1144 | 1181 | 39 |
| 1-Pentanol | dried hay | 1222 | 1255 | 32 |
| Octanal | fatty | 1256 | 1291 | 32 |
| (Z)-2-Heptenal | cooked fat | 1298 | 1319 | 54 |
| Nonanal | fatty, grease | 1381 | 1390 | 35 |
| 3,5-Octadien-2-ol | roasted vegetables | 1398 | ** | ** |
| 1,3-bis(1,1-dimethylethyl)-Benzene | oil | 1415 | 1426 | 55 |
| (E) 2-Octenal | rancid oil | 1423 | 1434 | 32 |
| 1-Octen-3-ol | grassy | 1449 | 1456 | 32 |
| Acetic Acid | vinegar | 1461 | 1465 | 34 |
| Decanal | rancid | 1501 | 1485 | 56 |
| Benzaldehyde | nutty, earthy | 1533 | 1508 | 24 |
| (E)-2-Nonenal | fried fat | 1541 | 1542 | 32 |
| (E)-2-Decenal | fatty | 1580 | ** | ** |
| d-Mannose | stale | 1662 | ** | ** |
| Benzeneacetaldehyde | floral, fruit | 1670 | 1646 | 32 |
| 1-(2-furyl)-1,2-Butanediol | sour cheese | 1694 | 1563 | 57 |
| 2,4-Dodecadialenal | fatty | 1729 | 2018 | 58 |
| 3-methyl-2(5H)-Furanone | cooked fat | 1750 | 1713 | 25 |
| 2-Undecenal | cheese | 1780 | 1755 | 52 |
| (E, E)-2,4-Decadienal | fatty | 1842 | 1819 | 32 |
| Hexanoic acid | sweaty, barnyard | 1887 | 1849 | 26 |
| 2-methyl-1-Hexadecanol | eggs | 1900 | ** | ** |
| Heptanoic acid | sweaty | 1991 | 1960 | 30 |
| 2-Pentadecanone | stale | 2046 | 2011 | 29 |
| Octanoic Acid | barnyard | 2084 | 2070 | 32 |
| Nonanoic acid | soapy | 2190 | 2169 | 32 |
| n-Decanoic acid | musty, stale | 2280 | 2279 | 26 |

**Table 4.** GC/MS-Olfactometry analysis of cultivated pork fat. Compounds were tentatively identified using NIST database matching; compound confirmation was made using published retention index values. Tentative identification is indicated by “**” (n=3).

| Compound | Aroma | RI (meas.) | RI (ref.) | Reference |
| --- | --- | --- | --- | --- |
| 2-ethyl-butanal | butter | 907 | 1018 | 46 |
| 3-methyl-3-buten-2-ol | sweet, caramel | 944 | ** | ** |
| Pentanal | sweet | 980 | 979 | 32 |
| Hexanal | sour, grassy | 1052 | 1078 | 39 |
| 2-pentylfuran | sour | 1177 | 1231 | 35 |
| Octanal | fatty, cooked fat | 1251 | 1291 | 32 |
| Dihydro-4,4-dimethyl-2(3H)-furanone | fatty | 1308 | ** | ** |
| Nonanal | fatty | 1353 | 1390 | 35 |
| 10-Methyl-8-tetradecen-1-ol acetate | butter, popcorn | 1392 | ** | ** |
| (E)-2-Octenal | nutty, cashews | 1406 | 1434 | 32 |
| 1-Octen-3-ol | musty | 1456 | 1456 | 32 |
| 10-Octadecenal | pork fat | 1466 | 1863 | 47 |
| 2-ethyl-1-Hexanol | potato, starch | 1491 | 1484 | 48 |
| Benzaldehyde | nuts | 1537 | 1508 | 24 |
| d-Mannitol | stale, barnyard | 1619 | ** | ** |
| (Z)-2-Decenal | cooked fat | 1634 | 1627 | 49 |
| Acetic acid | vinegar | 1645 | 1465 | 34 |
| Benzeneacetaldehyde | sour, fruit | 1668 | 1646 | 32 |
| Dimethyl Sulfoxide | stale | 1714 | 1582.3 | 37 |
| Pentanoic acid | cheese | 1735 | 1744 | 30 |
| 9-Octadecen-12-ynoic acid, methyl ester | cheese, sour | 1746 | 2478 | 50 |
| 2-methyl-hexanoic acid | cheese | 1749 | 1757 | 51 |
| 2-Undecenal | fruity, cheese | 1757 | 1755 | 52 |
| 2,4-Dodecadial | stale | 1837 | ** | ** |
| Hexanoic acid | barnyard, musty | 2054 | 1849 | 26 |
| Nonanoic acid | stale, musty | 2201 | 2169 | 32 |

## Discussion

The aroma profile of cell-cultivated adipose tissue was compared to that of conventional animal fat. The goal was to determine whether *in vitro*-grown adipose can be considered as an additive to enhance the organoleptic properties of alternative meat products based on aroma. To achieve this, we established a porcine DFAT clonal cell line produced by single cell seeding. These adipocytes were used to perform GC/MS and human sensory evaluation studies to analyze the volatile aroma compounds released during cooking. Additionally, we conducted an analysis using GC/MS-O to identify compounds associated with specific positive and negative aroma attributes that were identified by sensory evaluation panelists.

The majority of stable preadipocyte cell lines are from rodents, with few preadipocyte cell lines established for other species ^59^. To facilitate the study of pig fat for cellular agriculture, it is necessary to have access to an established adipogenic porcine cell line that can proliferate extensively and subsequently differentiate into mature adipocytes ^60^. A variety of porcine adipocyte progenitor cells have been screened for adipogenesis such as stromal vascular cells, pluripotent stem cells, and adipose derived stem cells ^23,38,61^. In our laboratory, the adipogenic potential of porcine stromal vascular cells (SVC) has been compared to that of DFAT cells (data not shown). The DFAT cells were capable of accumulating more lipids during adipogenesis ^23^. Therefore, porcine DFAT cells were used over SVCs for this project. Our previous work established a population of porcine DFAT cells which had a fatty acid composition closely resembling that of native pork fat. When grown in 2D, these cells could be aggregated into macroscale cultivated fat as a simplified method towards scalability. Using this method, we looked deeper into the utility of these cells for the formation of cultivated meat or more realistic meat alternatives.

We first established a porcine DFAT subclone from a previously isolated mixed population using single cell sorting and generated significant amounts of highly lipid laden adipocytes. Here, we show that among numerous successful single cell seedings, clone N47 demonstrated the highest intracellular lipid accumulation. However, N47 did not show more lipid accumulation than the original mixed population PDFAT. This was not expected, as previous studies showed that DFAT cells are often composed of more homogenous cell populations compared to stromal vascular and adipose derived stem cells due to their origin from highly homogenous fractions of mature adipocytes ^62–64^. It is likely that although N47 was selected as the highest preforming clone among approximately 576 clones, this screening was not large enough to uncover a clone with significantly higher lipid accumulation capacity than the mixed population.

As reported in our previous study, the mixed population PDFAT lipid morphology is multilocular and N47 exhibits the same multilocular phenotype representing an immature white adipocyte ^23^. Further culture optimization is required to achieve both higher lipid accumulation and unilocular lipids. In this study, optimization for PPARγ agonists dexamethasone, rosiglitazone and IBMX was conducted to improve lipid accumulation and promote a unilocular lipid phenotype. While limiting rosiglitazone to adipogenic induction media (first 2-3 days of adipogenesis) and lowering concentrations of IBMX to 0.1M increased lipid accumulation capacity, intracellular lipids remained multilocular. We assessed the influence of ascorbic acid in adipogenic differentiation medium related to adipogenesis ^65–69^. We found that ascorbic acid increased lipid accumulation in the cultivated porcine adipocytes. Additionally, we found that there was no significant difference in lipid accumulation when differentiating adipocytes in 10% and 1% serum. Thus, 10% FBS was used to differentiate adipocytes for GC/MS and sensory analysis to conserve yield, as serum has shown to increase cell adhesion. However, we did not find that the higher concentration of FBS prevented cell detachment throughout adipogenesis, therefore future studies may use 1% FBS to limit serum use ^70^. Ultimately, further optimization to mimic extensive lipid accumulation as achieved in native porcine adipocytes should be explored further to maximize the system described in the present work.

After generating the adipocytes, the cells were mechanically aggregated via cell scraping to generate fat tissue and subsequently used for GC/MS analysis. Compounds of interest included volatile and semi-volatile compounds that would be released during the cooking process of pork products. Specifically, for molecules associated with fatty aromas, total ion chromatograms revealed that conventional porcine fat samples contained many long and medium-chain fatty acids, the most abundant being oleic acid (9-octadecenoic acid), palmitic acid (n-hexadecanoic acid), capric acid (n-decanoic acid), myristic acid (tetradecanoic acid), palmitoleic acid (hexadec-9-enoic acid) and stearic acid (octadecenoic acid). Our results largely agree with the literature, as numerous groups have reported palmitic acid and stearic acid as the most prominent saturated fatty acids and oleic acid as the most prominent unsaturated fatty acid in porcine fat ^71–74^. Cultivated porcine fat samples contained mostly similar volatiles notably long chain fatty acids such as oleic acid, palmitic acid, and stearic acid. Other aroma attributes found in both samples such as “yeasty” and “cheesy” can be attributed to niacinamide (vitamin B3) and methyl ketones such as 2-Heptadecanone, 2-Heptanone and 2-Tridecanone, commonly detected in blue cheese ^75–78^. It should be noted that dimethyl sulfoxide was detected in cultivated fat samples among the top 20 volatiles, likely because of using DMSO as a solvent to deliver small molecules such as dexamethasone and IBMX in adipogenic media. Further research comparing the aroma profiles of native fats with cultured fats should seek alternative methods of incorporating hydrophobic compounds into the culture media to avoid any exogenous solvent additions like DMSO. For example, Intralipid uses lecithin as an emulsifier to deliver long-chain fatty acids and lecithin itself can be inductive of adipogenesis ^79,80^.

Molecules associated with aroma attributes such as “rancid” were also detected in both profiles. Fat samples contained pentanal, 2-pentylfuran, octanal, nonanal, and 1-octen-3-ol which correlate with oxidative sensory descriptors such as “rancid oil” in cooked meats ^81,82^. The cultivated fat indicated higher intensity peak areas for these factors. Additionally, cultivated fat contained oxidation related volatiles that were not found in conventional fat, such as heptanoic acid and 1-hexanol ^83,84^. The addition of ascorbic acid and other antioxidants has shown to reduce lipid oxidation and sulfur-containing compounds in pork products ^85,86^. Additionally, antioxidants may improve cell growth in serum-free conditions, which further emphasizes their addition when cultivating cells for food production ^87^. Other methods such as gamma (γ) irradiation and electron beam irradiation have been investigated as methods for the selective removal of rancid and off-odors from food products. Both are non-invasive and food safe for human consumption, as they do not impact the favorable aromas, nutritional qualities, or overall appearance and acceptability of the product ^88,89^. At high doses, γ irradiation may induce lipid oxidation and new off-odors, meaning further optimization of these methods is required to be applied to cultivated meat products ^89^.

Sensory evaluation with human subjects is a key method for studying the unique characteristics of food products and is largely unexplored for cultivated meat ^13^. Discrimination tasks are one of the simplest sensory analyses which require panelists to detect the difference between two samples. Here, we used discrimination testing to establish whether the aromas of cooked cultivated porcine fat could be distinguished from the aromas of cooked conventional porcine fat. The hypothesis tested was “with equivalent cooking parameters, are conventional and cultivated fat tissues perceptibly different products?”. Although the consumer panel was able to discriminate between the two samples, a significant preference for either fat sample was not found. A potentially confounding factor was the lack of extracellular matrix and scaffolding for the in vitro-grown fat compared to native fat tissue. There is currently no information on the specific contribution of ECM to the aroma and flavor of meat products. Additionally, these relatively immature adipocytes utilized here may contribute to a different aroma profile compared to native porcine fat that consists of mature fat cells. Metabolic characterization of differentiating adipocytes with GC/MS methods indicates that metabolic changes induced during stages of differentiation can be linked to the emission of different volatile and semi-volatile compounds ^90,91^. Lipid metabolism also shifts as adipogenic differentiation progresses, resulting in more long-chain fatty acids and desaturation in later stages ^90^. Therefore, aroma active volatiles at terminal stages of differentiation may have a different flavor profile than immature adipocytes. Additionally, in vitro-grown tissues were harvested and analyzed directly from culture whereas conventional meat samples underwent post-processing including carcass aging and long-term storage throughout the supply chain, which are known to significantly change the sensory characteristics of conventional meat ^92^. Future work may evaluate in vitro-grown pork fat after mimicking such conditions or conventional pork fat at different stages of production.

CATA profiling was conducted to further understand the differences in the aroma character of conventional and cultivated fat from the consumer perspective ^45^. Usage of attributes such as “fatty”, “savory”, “fried”, “meaty”, “bacon-like” and “roasted” were used equally by panelists to describe the two fat samples. Panelists used attributes “musty” and “barnyard” more often when describing cultivated porcine fat. As previously discussed, unlike cultivated porcine fat, commercial pork products undergo postmortem aging to increase the palatability attributes including aroma and flavor. This aging process leads to an increase in free amino acids and peptides which develops the “umami” flavor of pork via interactions with products of lipid oxidation such as peroxides. Additionally, products of lipid oxidation and nucleotide breakdown during this time are flavor precursors to the Maillard reaction, which contributes greatly to the browned, meaty aroma of cooked pork ^93,94^.

GC/MS-O was used to evaluate the range of both pleasant and foul aromas in the two tissue samples. “Pleasant” odors refer to aromas that led to an average increase in overall liking and “foul” odors refer to aromas that led to an average decrease in overall liking as determined by the sensory panelists. Twenty-two aroma-active compounds were found in cultivated pork fat while 26 were found in conventional pork fat. Similarly, previous studies found that with over 200 volatile compounds detected in beef, 36 were detected as key aroma compounds ^95^. Hexanoic acid, dimethyl sulfoxide, acetic acid and hexanal were detected in the top 20 volatiles from cultivated pork fat and appeared to also impact the sensory profile. Decanoic acid, 2-pentadecanone, 3-methyl-butanal, hexanoic acid, acetic acid and nonanal were detected in the top 20 volatiles from conventional pork fat and appeared to be sensory impactful. The sensory impact of DMSO in cultivated fat further supports the need to reduce the use of this agent in culture media and cryopreservation. With the exception of DMSO, these compounds have been previously reported in Yorkshire pork and beef samples ^84,96^.

Pleasant odors which occurred in both fat samples include fatty aldehydes such as pentanal, hexanal, octanal and nonanal. These compounds contributed to “sweet”, “buttery”, “fatty” and “greasy” aromas. Aldehydes are commonly known for their starchy, citrusy, fatty and waxy fragrance notes. In particular, C9 and C10 aldehydes such as nonanal and decanal (detected in cultivated porcine fat) are commonly found as skin odorants of various mammals resulting from the oxidation of sebaceous fatty acids ^97^. These saturated aldehydes are typical products of lipid oxidation and induce an increase of rancid oil flavors in high concentrations ^98^. However, these compounds presented a more positive, “fatty” aroma profile in the context of our pork fat samples. Foul odors detected in cultivated fat samples such as “barnyard”, “stale”, “musty” and “rancid oil” can be largely attributed to medium-chain fatty acid compounds such as pentanoic acid, hexanoic acid and heptanoic acid which are commonly created by the oxidation of the previously listed fatty aldehydes. These aldehydes along with furan compounds such as 2-pentylfuran and furanone found in both cultivated and conventional samples have been noted as the main components of pork flavor ^99^. They may also impact meat flavor by interacting with components of the Millard reaction. ^100^.

With future efforts, investigating the impact of storage on the volatile compounds of in vitro-grown fat may be useful to determine if chilled aging results in removal of negative aromas and a more similar volatile profile to conventional pork. Another proposed solution to these odors, instead of removing them completely, would be to reduce their intensity until they express a more pleasant aroma. Volatiles such as pentane which is present in cultivated fat can be attributed to having a “barnyard” odor. Pentane, among other volatiles, can change its sensory description as the concentration of the chemical compound increases. At low levels, the compound is “beany” while at high levels it is considered “sweaty”, and at its highest levels it has a “barnyard” odor. Similarly, hexanal, which is present at a higher concentration in our cultivated porcine fat than native porcine fat, presents a “pea pod” odor at low levels while at high levels presents “rancid”, “sour” and “oxidized oil” aromatics ^101–103^. Additionally, considering conventional pork fat contains many of the primary volatile markers for lipid oxidation due to either storage or cooking, reducing those in cultivated fat to equivalent levels should be sufficient. It’s important to note that even without these changes, consumers ultimately ranked cultivated and conventional porcine fat the same for overall liking.

### Conclusions

The primary objective of this paper was to address the following: What are the sensory characteristics of cell-cultivated adipose tissue, and how do these compare to traditional animal-derived fat? Aroma profiles of cell-cultivated adipose tissue were compared with conventional animal fat, aiming to evaluate in vitro-grown adipose as an additive to augment the organoleptic characteristics of alternative meat products. Through the establishment of a porcine DFAT clonal cell line and the application of advanced GC/MS techniques coupled with human sensory evaluations, we found that the volatile aroma compounds released during cooking were remarkably similar between the two fat sources. Despite some disparities in the presence and intensity of certain volatiles, the overall liking score from the sensory evaluation was similar for both in vitro and in vivo porcine adipose samples. The immature status of the adipocytes, the absence of extracellular matrix in the in vitro samples, the use of DMSO in the production process, and different post-harvest environments might be contributing factors to the differences in aroma profiles. Moving forward, our recommendations include refining the cell culture environment, investigating the role of extracellular matrix in aroma and flavor, and adjusting lipid supplementation during culture to better match the aroma profiles of conventional fat. As the field progresses, continued research is essential to not only closely replicate the aroma profile of native fats but also to ensure a complete and favorable sensory experience for consumers exploring alternative meats. Here we provided a methodological foundation to utilize and gain insight into these adipose sources. Importantly, the chemical-to-consumer links established by using this methodology can provide a guide to future studies on the topic. In addition, the data points to future opportunities to conduct molecular editing of flavors and odors to tailor outcomes, a major benefit to the processes involved in cultivated meat production.

## Methods

### Single Cell Seeding/Cell Line Development

Porcine DFAT cells were isolated from a 93-day-old female Yorkshire pig (DOB: 10/18/2021) ^23^. Single cell isolation from porcine DFAT cells was conducted at the Tufts University School of Medicine Flow Cytometry Core by fluorescence-activated cell sorting (FACS) with the FACSAria (BD Biosciences, Franklin Lakes, NJ, USA). Porcine DFAT cells were thawed at passage 3 and grown to 80% confluency in standard growth media containing Dulbecco’s Modified Eagle Medium (DMEM) + 20% Fetal Bovine Serum (FBS) + 100 μg/ml Primocin at 39°C and 5% CO_2_. DFAT cells were detached using Accumax (AM105; Innovative Cell Technologies Inc, San Diego, CA, USA) and resuspended in standard growth media. Cells were strained using a 40 µm strainer and diluted to achieve a suspension concentration of 1×10^6^ cells/mL. Subsequently, approximately 576 single cells were isolated into six 96-well plates. Single cells were seeded into wells containing 3 different culture media formulations: 1) The “Normal” group contained growth media (DMEM + 20% FBS + 100 μg/ml Primocin). 2) The “Conditioned” group contained 50% growth media and 50% conditioned media, which was growth media that had been incubated with DFAT cells for 2-3 days. 3) The “Vitronectin” group contained growth media supplemented with vitronectin peptide (VTN-N, 0.25 μg/cm^2^). Three days after seeding, wells were visually screened for the presence of single cells, or a single colony/cluster of several cells using an inverted light microscope (Olympus CKX53; Evident Life Science, Shinjuku-ku Tokyo, Japan). After eight days, cells were screened and VTN-N supplemented colonies were omitted from further screening due to poor proliferation. Wells in normal and conditioned media with successfully sorted cells were dissociated from the 96-well plates once a dense colony/cluster formed, to prevent the development of localized areas of cell confluence. After this initial passage, cells were cultured to 70-80% confluency and progressively moved to 24-well, 12-well, and 6-well plates. Cells were regularly inspected with a microscope for cell density and fat accumulation throughout six days of adipogenesis. Intracellular lipid accumulation of all viable clones was quantified with Oil Red O Stain Kit (ab150678; Abcam, Cambridge, UK) on day six (**Supplementary Fig. S1**). Five clones N18, N20, N47, C15, and C34 were selected based on previous proliferation screening and lipid accumulation screen (data not shown). Cells were monitored for nine passages before reducing the screen to clones N18, N47, and C34 based on lipid accumulation (data not shown). Clone N47 was finally selected based on a comparison of lipid accumulation for these top three clones as well as the mixed population PDFAT (**Supplementary Fig. S2**).

### Preadipocyte Cell Culture and Adipogenic Differentiation

Long term cell culture continued with PDFAT clone N47 grown using DMEM + 20% FBS + 100 µg/ml Primocin + 0.25 μg/cm^2^ laminin 511-E8 (N-892021; Iwai North America Inc., San Carlos, California, USA). Laminin 511-58 was used for ECM supplementation at every passage. The cells were cryopreserved in 90% culture medium and 10% dimethyl sulfoxide (DMSO). Cells were cultured in standard plasma treated tissue culture flasks and were handled gently during cell feedings to prevent detachment of adipocytes. During the aspiration of spent culture media, culture flasks were inverted and media was collected from the roof and sidewalls of the flasks. Additionally, new media was dispensed in the inverted position to avoid pipetting directly on the cells. The flasks were slowly returned to their upright positions to ensure gentle exposure of new media to the cells.

Porcine preadipocytes were grown until 100% confluency in culture medium. After remaining confluent for at least 24 hours, cells were switched to adipogenic induction media consisting of Advanced DMEM/F12 (12634-028; Thermo Fisher) with 10% FBS, 100 µg/ml Primocin, 500 µg/ml Intralipid (I141; Sigma), 0.1 mM 3-isobutyl-1-methylxanthine (IBMX, I5879; Sigma), 5 μM rosiglitazone (ROG, R0106; Thermo Fischer), 0.5 μM dexamethasone (DEX, AC23030; Thermo Fisher), 2 mM (1×) GlutaMAX (35050-061; Thermo Fisher), 20 µM biotin (B04631G; TCI America, Portland, OR, USA), and 10 µM calcium-D-pantothenate (P001225G; TCI America). Cells remained in adipogenic induction media for 2 days. During lipid accumulation, the same medium was used, omitting IBMX and ROG. Cells remained in the lipid accumulation media for 9-11 days. Media optimization was conducted for 5 μM rosiglitazone, 0.5 μM dexamethasone and 20ng/mL bone morphogenic protein-4 (BMP4, NBC119056; Thermo Fisher).

### Fat Harvest

After adipogenesis, the spent culture media was aspirated by gentle inversion of the flask as previously described. Adipocytes were rinsed with Dulbecco’s phosphate buffered saline (DPBS) 3-5 times and flasks were kept vertical for 1-2 mins to thoroughly drain DPBS and any remaining media. Once excess DPBS was aspirated, the adipocytes were harvested using a cell scraper (83.3952; Sarstedt, Nümbrecht, Germany) via raking motions from the front to the back of the flask. Periodically, additional liquid was gently aspirated from the flask without removing the scraped cells. The cell mass was pushed to the back of the flask and collected, then transferred into a pre-weighed 0.6 ml Eppendorf tube.

### Lipid Staining

Cultured adipocytes in 96-well tissue culture well plates were stained to confirm intracellular lipid accumulation. Here, PDFAT N47 adipocytes were stained using Oil Red O Stain Kit (ab150678; Abcam, Cambridge, UK). Cells were rinsed twice with DPBS and fixed with 4% paraformaldehyde (PFA) for 20 mins at room temperature. Oil Red O assessment was conducted according to the manufacturer’s protocol. Media and other liquids were aspirated with a pipette to avoid detachment of adipocytes. Cells were incubated with propylene glycol at RT for 5 min and Oil Red O was heated to 60°C. Propylene glycol was replaced by warmed Oil Red O and incubated at RT for 7 min. Cells were incubated with 85% propylene glycol in distilled water for 1 min and rinsed twice with distilled water. The results for Oil Red O staining were analyzed on a microplate reader at 515 nm.

### Native Porcine Adipose Samples

Pork fat tissue (*Sus scrofa domesticus*) from the belly was purchased commercially from a wholesale distributor. Samples were displayed at refrigerator temperatures in the store and transported to the laboratory at RT. The tissue was sliced, preserved with a food grade vacuum sealer and stored at −20°C. Subcutaneous fat from tissue samples was cut at room temperature and used for GC/MS and sensory evaluation experiments.

### GC/MS and GC/MS-Olfactory

The volatile profile of each tissue was collected using an Agilent 7890A gas chromatograph coupled with an Agilent 5977A mass selective detector (Agilent Technology, Santa Clara, CA, USA). The GC/MS was equipped with a robotic multi-purpose autosampler (MPS), dynamic headspace unit (DHS) thermal desorption unit (TDU), and cooled injection system (CIS4), each manufactured by Gerstel (Gerstel, Linthicum, MD). 50 mg of tissue was weighed into a 20 mL round bottom amber glass vial (Restek, Bellefonte, PA). The sampling protocol began with the MPS transferring the 20 mL vial into the incubation chamber of the DHS unit. The tube was first incubated at 120°C for 15 minutes; the vial was then purged with 200 mL of nitrogen at a flow of 100 ml/min; volatiles were trapped onto a Tenax filled TDU tube with 1,500 mL of nitrogen at a flow rate 100 mL/min at 120°C; the Tenax trap was then dried with 200 mL of nitrogen at 50 mL/min; lastly, the Tenax filled TDU tube was then transferred to the TDU for desorption. Prior to desorption, the CIS with glass bead liner, was cooled to −120°C with liquid nitrogen. Desorption was initiated by ramping the TDU from 40°C to 300°C at 720°C/min with a 5 min hold at 300°C. After desorption the TDU tube was removed from the TDU. The chromatographic analysis was initiated with the ramping of the CIS from −120°C to 280°C followed by a 5 min hold. The 7890A was outfitted with a DB-WAX UI capillary column (30 m, 0.25 mm i.d., 0.25 mm film thickness (Agilent Technologies, Santa Clara)). The 7890A oven temperature was programmed to hold at 40°C for 1 minute; then ramp at 10°C/min to 250°C, and then hold 250°C for 10 minutes. The helium carrier gas was set to a constant flow 1.2 mL/min. The MSD was set to a solvent delay for the initial 1.25 minutes, with an electron energy of −70 eV, a source temperature of 250°C, and quadrupole temperature of 150°C. Data was acquired in scan mode ranging from 35 *m/z* to 300 *m/z*. Triplicate injections of each sample were completed and unknown compounds were identified via matching MS fragmentation patterns against the NIST17 mass spectral libraries and published retention index values.

### GC/MS-O

Olfactory data was collected using an Agilent 7890A coupled with an Agilent 5975C mass selective detector (Agilent Technology, Santa Clara, CA, USA). The GC/MS was equipped with an MPS Robotic autosampler, DHS TDU, cooled injection system (CIS4), and olfactometer detection port (ODP3), each manufactured by Gerstel (Gerstel, Linthicum, MD). Tissue samples were processed using an equivalent DHS, TDU, and CIS settings as described for the GC/MS (see GC/MS and GC/MS-Olfactory). Chromatographic separation was achieved using an Agilent INNOWAX capillary column (30 m, 0.25 mm i.d., 0.25 mm film thickness (Agilent Technologies, Santa Clara)), with an equivalent oven temperature program.

### Sensory Panel Recruitment and Sample Preparation

A total of 55 participants were recruited from the students and staff in the Tufts School of Engineering, Medford, Massachusetts. Panelists were recruited via flier advertisements and email. Email databases included all undergraduate and graduate students in the Tufts School of Engineering. All participants received information regarding test type, product to be tested, and any risks or benefits of participating in the study. To participate, panelists had to be 18 years or older and willing to smell *in-vitro* and conventional porcine fat tissues. Prior to each test, panelists were provided with consent forms and given another brief description of the samples and testing method. The study assumed panelists were strictly untrained. Panelists who were considered untrained had no experience participating in either a triangle or check-all-that-apply sensory evaluation study. The entire study contained approximately 47% male, 47% female, and 6% non-binary identifying participants.

In-vitro and conventional porcine fat samples were thawed from frozen at −80°C and - 20°C samples, respectively. One-hundred mg samples were placed in headspace vials with the caps on and heated to 120°C in a commercial oven for 45 minutes. Separate samples were used for the triangle test and check-all-that-apply (CATA) analyses. Sample vials were warmed in 60°C bead baths throughout the duration of the sensory evaluation.

### Discrimination Task

To determine if subjects could discriminate between cultivated and conventional porcine fat samples, a triangle test was conducted. Data was collected in one day over a 6-hour period, during which five sessions with a maximum of twelve participants per session were tasked with differentiating between cultivated and conventional porcine fat aroma. A total of 55 participants completed the evaluation. Participants completed the session at an isolated desk with natural lighting. Samples of the adipose tissues were presented at random, making combinations such as AAB, ABA, BAA, BBA, BAB, and ABB. All six combinations of conventional and in-vitro porcine fat samples were presented. Vials containing samples were concealed to mask visual differences between fat tissues. Each triangle was presented in a series along with a cup of water. The presentation order of the six triangles was randomized for all participants. Samples were reheated at 60°C in a bead bath for three minutes between panelists.

### Consumer Preference

The 55 participants provided preference information through a questionnaire which used a 7-point hedonic scale and check-all-that-apply (CATA) task. A complete summary of consumer preference responses can be found as Supplementary Table S6.

Two fat samples were presented in random order for every participant. The ballot questions were as follows:

- Two questions using the 7-point hedonic scale ranging from Extremely Dislike to Extremely Like: “Please indicate your liking of the aroma found in Vial X.”
- Two check-all-that-apply (CATA) questions: “Which attributes would you use to describe the aroma found in Vial X? Check all that apply” from a list of the following descriptors: Meaty, Fresh Meat, Bacon-like, Fatty, Savory, Cheesy, Sweet, Fruity, Roasted, Rancid, Fried, Animal, Barnyard, Sweaty, Metallic, Cardboard, Sour, Musty.
- After samples were evaluated, panelists completed an exit survey for the following variables: “Age,” “Gender”, divided into four possible categories; “Income”, divided into nine possible categories; “Education”, six categories; “Would you be willing to buy cultivated meat if it has the same price as conventional meat?”, Yes or No; Would you be willing to pay 10% more?”, Yes or No; Would you be willing to buy it with a 10% discount?”, Yes or No; “What do you like about the idea of cultivated meat?”, “What do you dislike about the idea of cultivated meat?”

### Statistical Analysis

Statistical analyses were conducted GraphPad Prism 10.1.1 and R version 4.3.2. The specific analyses include Analysis of variance (ANOVA) with Tukey’s post-hoc tests and Fisher’s exact tests. Error bars and ± ranges represent standard deviations. A p-value of 0.05 was used for statistical significance. All experiments in this study were carried out with at least triplicate samples (n≥3).

### Ethics Declaration

Isolation of adipocyte progenitor cells from a 93-day-old female Yorkshire pig (*Sus domesticus)* was approved by the Tufts University Medical Center and sourced from the Tufts Comparative Medicine Services (CMS). CMS methods are in accordance with the guidelines of the United States Department of Agriculture (USDA), Office of Laboratory Animal Welfare (OLAW), Massachusetts Department of Public Health (MDPH) and Association for Assessment and Accreditation of Laboratory Animal Care International (AAALAC). Additionally, all methods were conducted in accordance with ARRIVE guidelines 2.0.

The sensory evaluation study was approved by the Tufts University Health Sciences Institutional Review Board (IRB) for research involving human subjects (Study 00003031). All experiments were performed in accordance with the Tufts University Health Sciences IRB guidelines for exempt studies. Informed consent was obtained from all participants.

## Acknowledgements

We thank the United States Department of Agriculture (2021-69012-35978) and the Tufts University Center for Cellular Agriculture Consortium (TUCCA-C). We are grateful to Stephen Kwok, Allen Parmelee and Albert Tai from the Tufts University Flow Cytometry Core for assistance with FACS. We also thank Sean Cash from the Tufts Friedman School of Nutrition for assistance with the sensory evaluation consumer acceptance questionnaire. Figures were created using Biorender.com and Adobe Illustrator version 28.2.

## Contributions

E.T.L. contributed to methodology, conducting experiments, preforming data analysis, designing figures, and writing the main manuscript text. J.S.K.Y.Jr. contributed to methodology, conducting experiments, and drafting manuscript text. K.L.Z. contributed to conducting experiments and drafting manuscript text. K.F. contributed to methodology. S.C.F. contributed to methodology, conducting experiments, and designing figures. D.L.K. contributed to experimental design and manuscript drafting. All the authors have read and contributed to manuscript editing.

## Corresponding Author

Correspondence to David L. Kaplan

## Data Availability

The GC/MS datasets generated during and/or analyzed during the current study are available from the corresponding author on request.

