## Supplementary Information for "Chemical and Sensory Analyses of Cultivated Pork Fat Tissue as a Flavor Enhancer for Meat Alternatives"

### Supplementary Figure S1: Clonal isolation adipogenesis screening

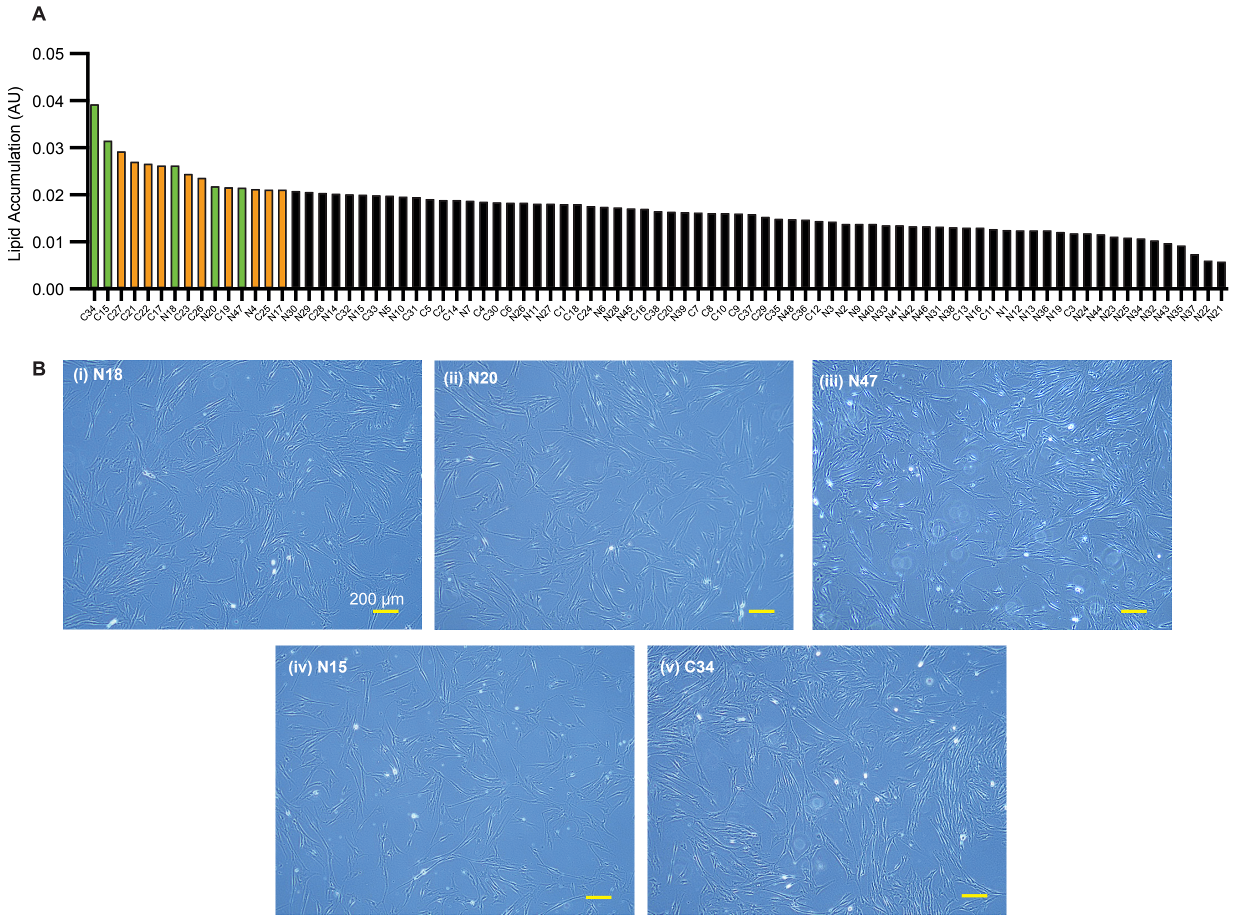


**Supplementary Figure S1.** Screening of clones isolated from mixed population PDFAT. (**A**) Lipid accumulation screen of normal and conditioned media clones. Clones N1-N48 and C1-C38 were stained with Oil Red O. All clones in vitronectin supplemented media were omitted due to poor proliferation. Clone N8 was omitted due to absorbance value 0. Top 15 clones are highlighted (orange) and clones picked for further screening are highlighted (green). Five clones were selected based on lipid accumulation and previous proliferation screening. (**B**) Morphology of top five clonal populations during proliferation (**i**) N18 (**ii**) N20 (**iii**) N47 (**iv**) C15 (**v**) C34 (passage 3). Scale bar represents 200 µm for all images. Scale bar represents 200 µm for all images.

**Supplementary Figure S2: Screening of clones N18, N47 and C34**

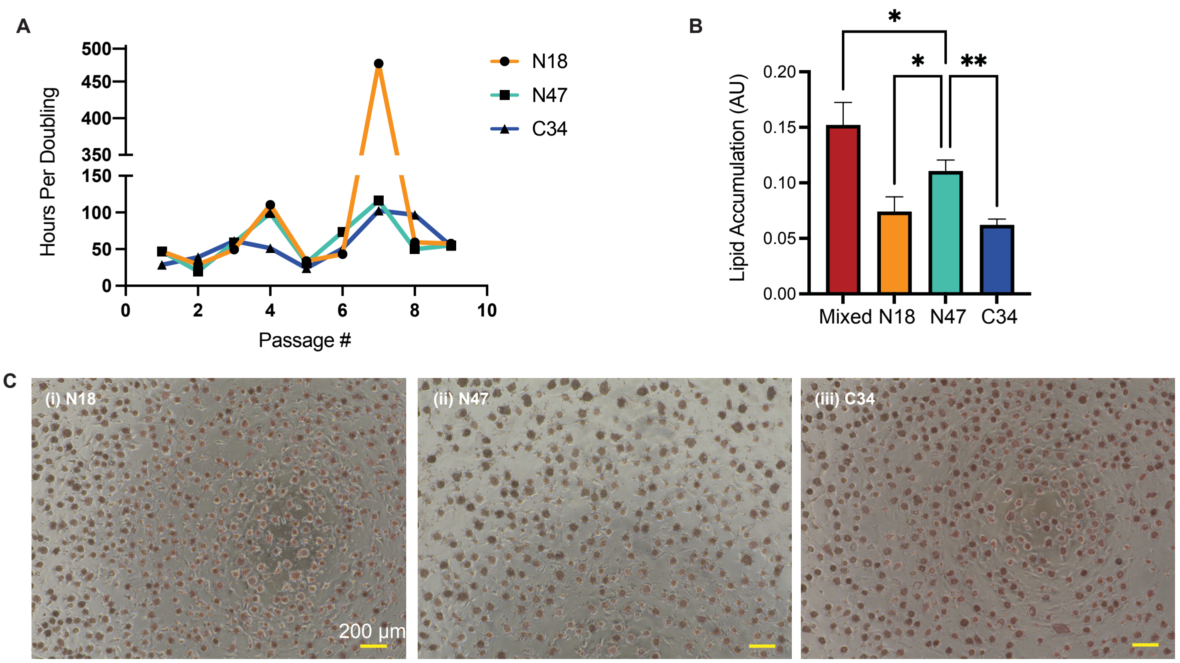


**Supplementary Figure S2.** Screening of clones N18, N47 and C34. (**A**) Hours per doubling (doubling time) of top three clones N18, N47 and C34 up to passage 9 (**B**) Lipid accumulation screen of mixed population PDFAT and top three clones (P8) determined by proliferation speed up to P6. Screening continued with clone N47. (**C**) Clones (**i**) N18 (**ii**) N47 (**iii**) C34 after six days of adipogenesis and stained with Oil Red O. Scale bar represents 200 µm for all images.

### Supplementary Figure S3: PDFAT lipid morphology

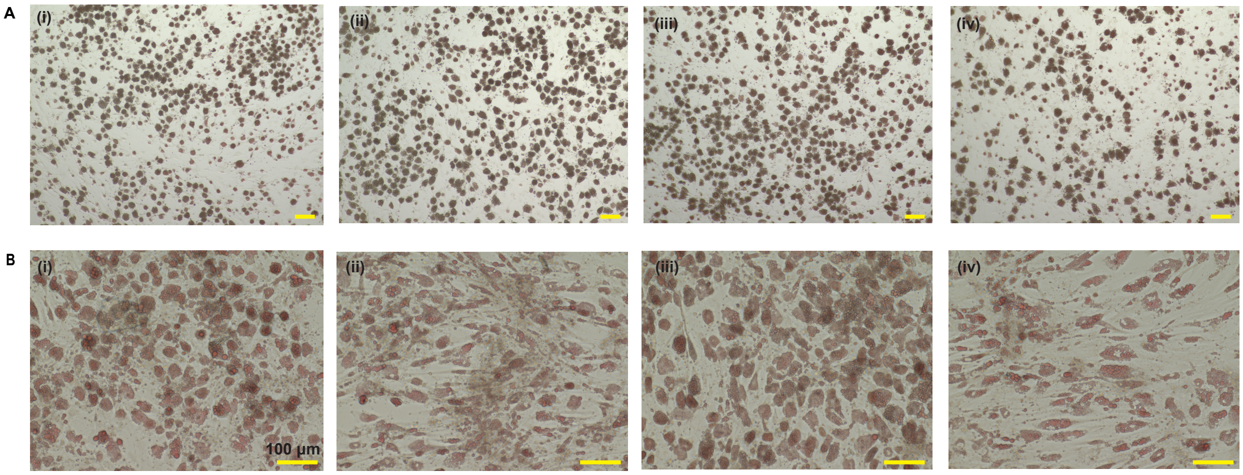


**Supplementary Figure S3.** Lipid morphology of N47 clones and PDFAT mixed population throughout IBMX, dexamethasone and rosiglitazone media optimization efforts. (**A)** Morphology of N47 cells with various concentrations of IBMX present in the induction media (P12) (A,**i**) 0 mM (A,**ii**) 0.1 mM (A,ii**i**) 0.25 mM (A,**iv**) 0.5 mM. Scale bar represents 100 µm (**B**) Morphology of mixed population PDFAT (B,**i**) +DEX +ROG (**B,ii**) –DEX +ROG (**B,iii**) +DEX –ROG (**B,iv**) –DEX –ROG in accumulation media (P3). Scale bar represents 100 µm.

**Supplementary Figure S4: N47 media optimization**

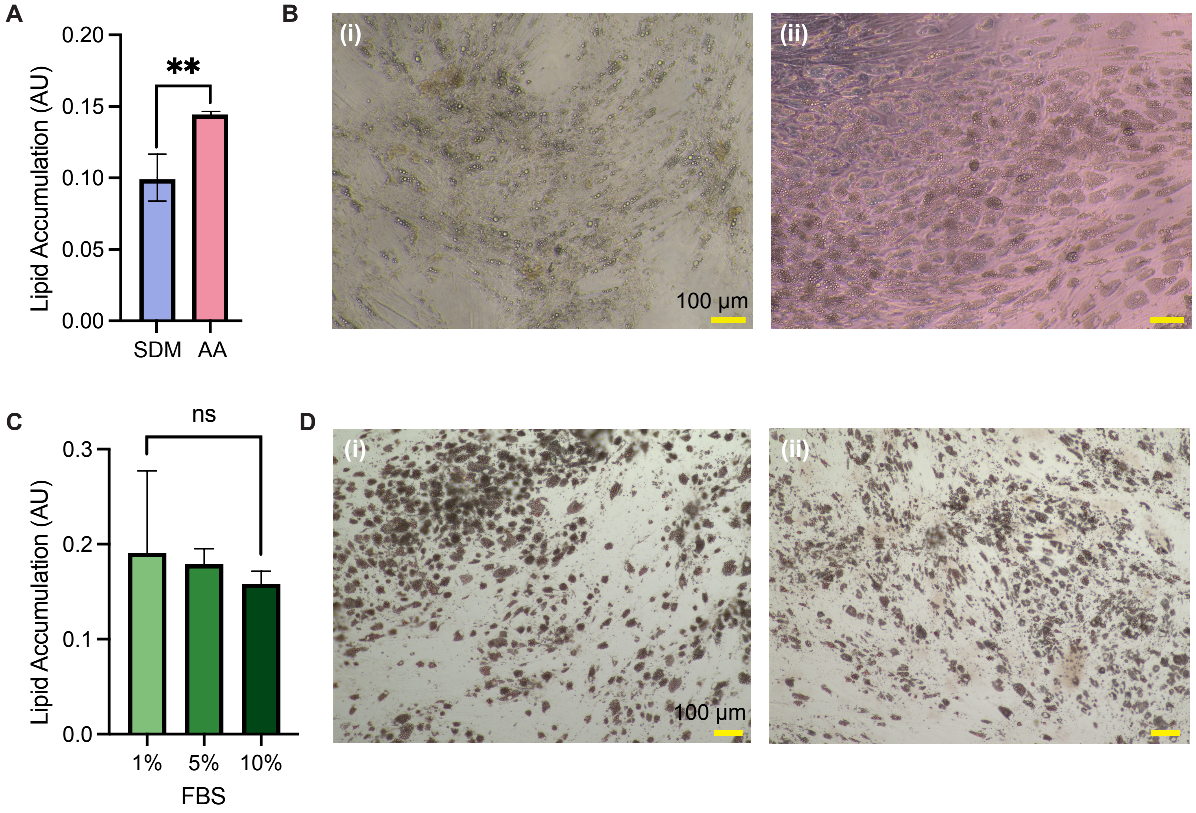


**Supplementary Figure S4.** Ascorbic acid adipogenic media screen for clone N47. (**A**) Lipid accumulation screen where SDM= standard growth media and AA = SDM supplemented with 113 µM ascorbic acid as described in Jurek et al. (n= 3). (**B**) N47 (passage 9) after nine days of adipogenesis with (**i**) SDM and (**ii**) AA. Scale bar represents 100 µm (n=4). **(C)** FBS concentration in induction and lipid accumulation medias with clone N47 (passage 23). (**D**) N47 (passage 23) after 10 days of adipogenesis with (**i**) 1% FBS and (**ii**) 10% FBS. Scale bar represents 100 µm (n=3).

**Supplementary Table S1: Total compound comparison of conventional and cultivated porcine fat.**

(**) tentatively identified with NIST hit.

| Compound | Peak Area | | Retention Time (mins.) | | RI (meas.) | |
|---|---|---|---|---|---|---|
| | In vitro | In vivo | In vitro | In vivo | In vitro | In vivo |
| 1-Dodecanol | 0.419 | 0.734 | 15.815 | 15.577 | 1995 | 1972 |
| 1-Hexadecanol | 1.171 | 0.382 | 20.115 | 20.441 | 2464 | 2503 |
| 1-Octen-3-ol | 0.177 | 0.472 | 9.579 | 9.578 | 1452 | 1452 |
| 1-Pentanol | 0.537 | 0.499 | 6.935 | 6.938 | 1257 | 1258 |
| 1-Tetradecanol | 0.379 | 0.117 | 17.584 | 17.576 | 2178 | 2177 |
| 2-Heptadecanone | 0.336 | 1.329 | 18.124 | 18.128 | 2236 | 2237 |
| 2-Octenal, (E)- | 0.426 | 0.539 | 9.340 | 9.337 | 1434 | 1434 |
| 2-Propenoic acid, 2-methyl- | 0.145 | 0.269 | 12.493 | 12.496 | 1688 | 1688 |
| 2,2,4-Trimethyl-1,3-pentanediol diisobutyrate | 0.619 | 1.568 | 14.699 | 14.695 | 1887 | 1887 |
| 2(3H)-Furanone, 5-ethyldihydro- | 0.420 | 0.177 | 12.720 | 2.208 | 1707 | 698 |
| 2(5H)-Furanone, 3-methyl- | 0.313 | 0.555 | 12.868 | 12.870 | 1721 | 1721 |
| 2H-Pyran-2-one, tetrahydro-6-nonyl- | 0.336 | 0.476 | 21.799 | 21.797 | 2672 | 2672 |
| 2H-Pyran-2-one, tetrahydro-6-pentyl- | 0.543 | 1.191 | 17.844 | 17.843 | 2205 | 2205 |
| 2H-Pyran-2-one, tetrahydro-6-undecyl- | 1.020 | 1.051 | 23.826 | 23.826 | 2901 | 2901 |
| 3-Methyl-hexanoic acid | 0.433 | 0.443 | 15.369 | 15.352 | 1952 | 1951 |
| 3-Nonen-2-one | 0.873 | 0.405 | 14.581 | 16.670 | 1877 | 2082 |
| 4H-Pyran-4-one, 3,5-dihydroxy-2-methyl- | 0.572 | 0.251 | 18.604 | 18.625 | 2289 | 2291 |
| 5-Ethylcyclopent-1-enecarboxaldehyde | 0.150 | 0.158 | 9.191 | 9.197 | 1422 | 1422 |
| 7,9-Di-tert-butyl-1-oxaspiro(4,5)deca-6,9-diene-2,8-dione | 0.373 | 0.403 | 22.354 | 22.354 | 2741 | 2741 |
| 9-Octadecenoic acid, (E)- | 4.546 | 19.700 | 27.141 | 27.148 | 3066 | 3066 |
| Acetic acid | 3.784 | 1.612 | 10.416 | 10.428 | 1517 | 1518 |
| Acetic acid, methyl ester | 0.323 | 0.704 | 7.595 | 7.602 | 1302 | 1303 |
| Acetoin | 1.343 | 0.280 | 7.416 | 7.418 | 1290 | 1290 |
| Acetophenone | 0.196 | 0.164 | 12.092 | 12.100 | 1655 | 1655 |
| Benzaldehyde | 0.175 | 0.420 | 10.522 | 10.523 | 1525 | 1526 |

| | | | | | | |
|---|---|---|---|---|---|---|
| Benzoic acid | 0.696 | 0.532 | 19.950 | 19.923 | 2445 | 2441 |
| Benzyl alcohol | 0.200 | 0.141 | 14.607 | 14.614 | 1879 | 1880 |
| Butanal, 3-methyl- | 4.867 | 2.504 | 2.903 | 2.907 | 852 | 852 |
| Butanoic acid | 0.506 | 0.817 | 11.761 | 11.755 | 1626 | 1626 |
| Butyrolactone | 0.579 | 0.356 | 11.840 | 11.835 | 1633 | 1633 |
| Cyclodecane | 0.518 | 0.382 | 15.575 | 15.577 | 1972 | 1972 |
| Delta Dodecalactone | 0.552 | 1.067 | 19.904 | 19.907 | 2439 | 2439 |
| Dibutyl phthalate | 1.051 | 0.448 | 22.046 | 22.044 | 2703 | 2703 |
| Dimethyl ether | 0.244 | 0.537 | 3.082 | 3.078 | 864 | 863 |
| Dodecanoic acid | 0.918 | 5.855 | 20.271 | 20.277 | 2483 | 2483 |
| Dodecyl acrylate | 0.226 | 0.168 | 15.824 | 15.828 | 1996 | 1996 |
| Formic acid, octyl ester | 0.272 | 0.324 | 10.975 | 10.974 | 1562 | 1562 |
| Furan, 2-pentyl- | 3.761 | 0.674 | 6.609 | 6.595 | 1234 | 1233 |
| Hexadecane | 1.163 | 0.296 | 11.457 | 11.448 | 1600 | 1599 |
| Hexadecanoic acid, butyl ester | 0.324 | 0.121 | 19.903 | 19.898 | 2439 | 2438 |
| Hexanal | 3.141 | 0.948 | 4.673 | 4.627 | 1093 | 1089 |
| Hexanoic acid | 5.139 | 2.231 | 14.229 | 14.224 | 1844 | 1844 |
| n-Hexadecanoic acid | 8.010 | 20.138 | 23.886 | 23.900 | 2907 | 2908 |
| Naphthalene-D8 | 1.000 | 1.000 | 13.147 | 13.144 | 1746 | 1746 |
| Niacinamide | 3.801 | 2.637 | 24.614 | 24.603 | 2972 | 2971 |
| Nonanal | 0.734 | 1.572 | 8.855 | 8.849 | 1396 | 1395 |
| Nonanoic acid | 1.072 | 1.371 | 17.464 | 17.462 | 2165 | 2165 |
| Octadecanoic acid | 2.827 | 6.738 | 26.613 | 26.617 | | |
| Octanal | 0.577 | 0.770 | 7.432 | 7.426 | 1291 | 1291 |
| Octanoic acid | 0.919 | 6.353 | 16.438 | 16.434 | 2059 | 2058 |
| Palmitoleic acid | 1.203 | 10.155 | 24.293 | 24.297 | 2943 | 2944 |
| Pentadecanoic acid | 0.372 | 0.471 | 22.855 | 22.858 | 2800 | 2801 |
| Pentanal | 0.932 | 0.482 | 3.469 | 3.472 | 886 | 887 |
| Pentanoic acid | 0.684 | 0.193 | 13.041 | 13.048 | 1737 | 1737 |
| Phthalic acid, isobutyl 4-octyl ester | 1.121 | 0.500 | 20.806 | 20.803 | 2548 | 2547 |
| Propanal, 2-methyl- | 1.570 | 0.183 | 2.261 | 2.225 | 804 | 800 |
| Propanoic acid | 1.407 | 0.278 | 10.666 | 10.663 | 1537 | 1537 |
| Styrene | 0.194 | 0.570 | 6.964 | 6.972 | 1260 | 1260 |
| Tetradecane | 2.223 | 0.271 | 8.923 | 8.904 | 1400 | 1399 |
| Tetradecanoic acid | 1.911 | 11.990 | 21.981 | 21.986 | 2695 | 2696 |

**Supplementary Table S2. Compounds unique to cultivated porcine fat samples**
(**) tentatively identified with NIST hit.

| Compound | Peak Area | Retention Time (mins.) | RI (meas.) |
|---|---|---|---|
| 2-Oxetanone, 4-methyl- | 9.538 | 2.910 | 852 |
| 2-Heptanone | 4.503 | 5.979 | 1189 |
| Dimethyl Sulfoxide | 4.349 | 11.285 | 1587 |
| 3-Pentanamine | 3.444 | 2.889 | 851 |
| 2-Decanone | 3.100 | 10.173 | 1497 |
| Cyclobutanol | 3.094 | 2.899 | 852 |
| 2-Nonanone | 2.962 | 8.809 | 1392 |
| Phthalic acid, di(2-propylpentyl) ester | 2.216 | 27.486 | ** |
| 2(3H)-Furanone, dihydro-5-propyl- | 1.830 | 16.236 | 2038 |
| 5-Cyclohexyl-1-pentene | 1.623 | 28.099 | |
| trans-3-Nonen-2-one | 1.589 | 16.683 | 2083 |
| Ethanone, 1-(3-butyloxiranyl)- | 1.477 | 11.948 | 1643 |
| Methane, isocyanato- | 1.392 | 2.877 | 850 |
| 1-Propanamine, 2-methyl-N-(3-methylbutylidene)- | 1.195 | 4.365 | 1065 |
| 4H-Pyran-4-one, 2,3-dihydro-3,5-dihydroxy-6-methyl- | 1.136 | 18.452 | 2272 |
| (E)-9-Octadecenoic acid ethyl ester | 1.076 | 20.294 | 2485 |
| n-Pentadecanol | 1.034 | 18.522 | 2280 |
| Dodecane | 0.918 | 6.132 | 1198 |
| 2(3H)-Furanone, 5-dodecyldihydro- | 0.841 | 23.258 | 2843 |
| 2(3H)-Furanone, 5-butyldihydro- | 0.800 | 15.838 | 1997 |
| Heptanoic acid | 0.796 | 15.357 | 1951 |
| Gamma tetradecalactone | 0.774 | 21.359 | 2615 |
| Oleic Acid | 0.680 | 27.126 | |
| Oxazole | 0.650 | 3.901 | 1019 |
| 2(3H)-Furanone, dihydro-3-hydroxy-4,4-dimethyl-, (.+/-.)- | 0.648 | 16.182 | 2033 |
| Phthalic acid, di(6-methylhept-2-yl) ester | 0.643 | 27.460 | ** |
| 2,6,10,14-Hexadecatetraen-1-ol, 3,7,11,15-tetramethyl-, acetate, (E,E,E)- | 0.642 | 20.725 | 2538 |
| Hexadecanoic acid, ethyl ester | 0.638 | 18.322 | 2258 |
| Farnesyl butanoate | 0.619 | 20.735 | 2539 |
| (Z)-9-octadecen-4-olide | 0.610 | 26.359 | ** |
| 1-Butanamine, N-butylidene- | 0.562 | 6.192 | 1203 |
| Undecane, 3,8-dimethyl- | 0.553 | 13.766 | 1801 |
| 9,12-Octadecadienoic acid (Z,Z)- | 0.550 | 20.448 | 2504 |

| | | | |
|---|---|---|---|
| Methyl 3-oxohexadecanoate | 0.549 | 16.110 | 2025 |
| 2-Fluoroaniline, N-pentyl- | 0.531 | 19.523 | 2394 |
| 3-Tetradecanol | 0.529 | 15.846 | 1998 |
| 1,3-Butadiyne, 1,4-difluoro- | 0.508 | 2.892 | 851 |
| Piperidine, 1-ethyl- | 0.503 | 7.642 | 1306 |
| 1-Butene, 3-butoxy-2-methyl- | 0.486 | 19.303 | 2369 |
| 6-Undecanol | 0.483 | 23.681 | 2887 |
| 2,6-Di-tert-butyl-4-hydroxy-4-methylcyclohexa-2,5-dien-1-one | 0.483 | 16.840 | 2099 |
| 4-Oxohex-2-enal | 0.479 | 13.342 | 1764 |
| 2(3H)-Furanone, dihydro-5-(2-octenyl)-, (Z)- | 0.472 | 19.651 | 2409 |
| Butanoic acid, 2-hexenyl ester, (E)- | 0.463 | 18.389 | 2265 |
| 1H-Benzocycloheptene, 4,4a,5,6,7,8,9,9a-octahydro-4a-methyl-, trans- | 0.458 | 21.745 | 2665 |
| Benzene, 1,1'-(1,2-cyclobutanediyl)bis-, cis- | 0.447 | 19.680 | 2412 |
| Heptadecane | 0.445 | 12.631 | 1699 |
| 2-Dodecanol, 2-methyl- | 0.442 | 15.848 | 1998 |
| Cyclohexanone, 2-propyl- | 0.440 | 15.280 | 1944 |
| 1-Octadecanol | 0.434 | 21.153 | 2590 |
| Pentanoic acid, 5-hydroxy-, 2,4-di-t-butylphenyl esters | 0.426 | 18.759 | 2306 |
| 3-Octen-2-one | 0.421 | 9.058 | 1411 |
| Octadecanoic acid, ethyl ester | 0.415 | 20.116 | 2464 |
| 1-Butanamine, 2-methyl-N-(2-methylbutylidene)- | 0.406 | 5.369 | 1147 |
| 2-Pentadecanone, 6,10,14-trimethyl- | 0.401 | 17.130 | 2130 |
| 1-Butanamine, 3-methyl-N-(3-methylbutylidene)- | 0.400 | 6.433 | 1221 |
| Benzeneacetic acid | 0.384 | 20.926 | 2562 |
| Piperidine, 1-(1-methylpentyl)- | 0.366 | 15.217 | 1937 |
| 2-Pyrrolidinone | 0.364 | 16.337 | 2048 |
| 2-Octanone | 0.361 | 7.385 | 1288 |
| Hexadecenoic acid, Z-11- | 0.360 | 24.312 | 2945 |
| Acetamide | 0.360 | 13.330 | 1763 |
| Undecane, 2,9-dimethyl- | 0.358 | 10.216 | 1500 |
| Cyclohexanone | 0.358 | 15.292 | 1945 |
| 2-Octenoic acid | 0.350 | 17.637 | 2183 |
| Cyclohexanone, 2-butyl- | 0.348 | 13.972 | 1820 |
| Amantadine | 0.348 | 11.701 | 1621 |
| 2-Pentanol, 2,4-dimethyl- | 0.337 | 21.069 | 2579 |
| Pyridine, 3-phenyl- | 0.333 | 18.299 | 2255 |
| Isoxazole | 0.326 | 4.023 | 1032 |

| | | | |
|---|---|---|---|
| (Z)-10-Pentadecene-1-ol | 0.323 | 18.791 | 2310 |
| 2-Octene, 4-ethyl-, (E)- | 0.322 | 10.702 | 1540 |
| Trifluoroacetic acid,n-tridecyl ester | 0.298 | 16.600 | 2075 |
| 2-Tridecanol | 0.289 | 17.094 | 2126 |
| 4,8,12,16-Tetramethylheptadecan-4-olide | 0.282 | 24.696 | 2979 |
| 1-Naphthalenepropanol, .alpha.-ethenyldecahydro-.alpha.,5,5,8a-tetramethyl-2-methylene-, [1S-[1.alpha.(R*),4a.beta.,8a.alpha.]]- | 0.281 | 21.862 | 2680 |
| (Z)-Dodec-5-en-4-olide | 0.281 | 21.528 | 2637 |
| 3,7-Dimethyl-1-octyl methylphosphonofluoridate | 0.264 | 11.890 | 1638 |
| 1-Tricosene | 0.248 | 13.175 | 1749 |
| Pentane, 1-(2-butenyloxy)-, (E)- | 0.247 | 11.883 | 1637 |
| Tetrahydrofuran-2-one, 5-[1-hydroxyhexyl]- | 0.246 | 22.546 | 2764 |
| 2-Tetradecanol | 0.239 | 16.107 | 2025 |
| Aziridine, 1-ethenyl- | 0.231 | 3.887 | 1018 |
| 5-Thiazoleethanol, 4-methyl- | 0.225 | 18.829 | 2314 |
| Homosalate | 0.224 | 20.132 | 2466 |
| Pentanoic acid, 2-methyl-, anhydride | 0.222 | 7.872 | 1324 |
| Phenol, o-amino- | 0.218 | 18.661 | 2295 |
| 1-Hexanol, 2-ethyl- | 0.217 | 10.108 | 1492 |
| Isopropylcyclobutane | 0.214 | 9.671 | 1459 |
| Methanamine, N-pentylidene- | 0.208 | 4.286 | 1058 |
| Undecane, 4,4-dimethyl- | 0.208 | 10.779 | 1547 |
| Isopropyl myristate | 0.205 | 16.257 | 2040 |
| 2H-Pyran-2-one, tetrahydro- | 0.200 | 13.896 | 1813 |
| 3-Acetamidofuran | 0.200 | 19.956 | 2445 |
| Succinimide | 0.196 | 20.177 | 2472 |
| (2E,6E,10E)-3,7,11,15-Tetramethylhexadeca-2,6,10,14-tetraen-1-yl formate | 0.192 | 20.727 | 2538 |
| 1,2(axial)-Dimethyl-trans-decahydroquinol-4-one | 0.191 | 9.869 | 1474 |
| 2-n-Butyl furan | 0.186 | 5.229 | 1137 |
| 3-Ethylcyclopentanone | 0.182 | 8.054 | 1338 |
| Hexanoic acid, anhydride | 0.181 | 13.373 | 1766 |
| 2-Octenal, 2-butyl- | 0.181 | 12.309 | 1673 |
| 1-Hexanol | 0.179 | 8.315 | 1357 |
| Phytol | 0.177 | 21.366 | 2616 |
| Hexanamide | 0.175 | 16.928 | 2108 |
| Hexanedioic acid, dioctyl ester | 0.175 | 23.155 | 2832 |
| Benzophenone | 0.167 | 20.352 | 2492 |

| | | | |
|---|---|---|---|
| 2,2-Diethyl-N-ethylpyrrolidine | 0.163 | 18.593 | 2287 |
| Benzamide, N,N-diethyl-4-methyl- | 0.162 | 18.673 | 2296 |
| N-Benzoyl-L-proline | 0.160 | 21.827 | 2676 |
| 1-Dodecanol, 3,7,11-trimethyl- | 0.155 | 13.688 | 1794 |
| Phenol | 0.154 | 15.877 | 2001 |
| Decane, 5-propyl- | 0.148 | 7.544 | 1298 |

**Supplementary Table S3. Compounds unique to conventional pork fat**
(**) tentatively identified with NIST hit.

| Compound | Peak Area | Retention Time (mins.) | RI (meas.) |
|---|---|---|---|
| n-Decanoic acid | 15.507 | 18.452 | 2272 |
| 2,6-Dimethylbicyclo[3.2.1]octane | 9.191 | 28.104 | ** |
| 9(E),12(E)-Conjugated linoleic acid | 8.319 | 28.113 | ** |
| 2-Pentadecanone | 7.523 | 16.125 | 2027 |
| Glycerin | 7.168 | 18.850 | 2316 |
| 9-Decenoic acid | 3.970 | 18.982 | 2332 |
| Silane, diethoxydimethyl- | 1.527 | 12.418 | 1682 |
| 2,4-Decadienal | 1.526 | 13.913 | 1815 |
| Propanoic acid, 2-methyl-, 3-hydroxy-2,2,4-trimethylpentyl ester | 1.383 | 14.544 | 1873 |
| 1-Methyl-5-fluorouracil | 1.351 | 18.457 | 2273 |
| Propanoic acid, 2-methyl-, 2-ethyl-3-hydroxyhexyl ester | 1.205 | 14.758 | 1893 |
| Oxirane, trimethyl- | 1.189 | 2.890 | 851 |
| 2-Heptenal, (E)- | 0.953 | 7.922 | 1328 |
| 2-Tridecanone | 0.952 | 13.898 | 1813 |
| Benzeneacetaldehyde | 0.856 | 11.953 | 1643 |
| 2-Decenal, (E)- | 0.831 | 12.012 | 1648 |
| Thiophene, 2,3-dihydro- | 0.818 | 19.436 | 2384 |
| 2-Undecenal | 0.793 | 13.264 | 1757 |
| Butane,1,2,4-trichloro-heptafluoro- | 0.751 | 12.453 | 1685 |
| cis-10-Heptadecenoic acid | 0.748 | 25.562 | ** |
| Butanoic acid, 3-methyl- | 0.672 | 12.257 | 1669 |
| 2,4-Decadienal, (E,E)- | 0.661 | 13.389 | 1768 |
| 2-Nonenal, (E)- | 0.640 | 10.693 | 1540 |
| Butanimidamide | 0.616 | 2.901 | 852 |
| 2(3H)-Furanone, 5-hexyldihydro- | 0.594 | 19.448 | 2385 |
| Heptanal | 0.591 | 5.984 | 1189 |
| Myristoleic acid | 0.578 | 22.353 | 2741 |
| Oxacyclopentadecan-2-one | 0.577 | 22.295 | 2734 |
| 2(3H)-Furanone, 5-acetyldihydro- | 0.528 | 16.482 | 2063 |
| Vinyl crotonate | 0.485 | 12.856 | 1720 |
| 2,5-Dimethylfuran-3,4(2H,5H)-dione | 0.471 | 16.196 | 2034 |
| Oxacyclotridecan-2-one | 0.470 | 20.779 | 2544 |
| p-Cresol | 0.462 | 16.635 | 2079 |
| Dodecanal | 0.457 | 12.770 | 1712 |

| | | | |
|---|---|---|---|
| Cyclotetradecane | 0.452 | 19.398 | 2379 |
| 3(2H)-Furanone, 4-hydroxy-5-methyl- | 0.438 | 17.014 | 2118 |
| Ethanol, 2-(hexyloxy)- | 0.422 | 11.644 | 1616 |
| 2,4-Imidazolidinedione, 1-methyl- | 0.407 | 22.169 | 2718 |
| Benzene, 1,3-bis(1,1-dimethylethyl)- | 0.396 | 9.303 | 1431 |
| Z-7-Tetradecenoic acid | 0.394 | 22.364 | 2742 |
| Indole | 0.392 | 19.967 | 2447 |
| 2,4-Decadienal, (E,Z)- | 0.392 | 13.400 | 1769 |
| Pentadecane | 0.375 | 10.208 | 1499 |
| Phenol, 4-(1-methylpropyl)- | 0.360 | 14.314 | 1852 |
| Ethanol, 2-(2-butoxyethoxy)- | 0.359 | 13.763 | 1800 |
| 1,4-Dioxane-2,5-dione, 3,6-dimethyl- | 0.352 | 16.197 | 2034 |
| 1,4-Dioxane-2,5-dione, 3,6-dimethyl-, (3S-cis)- | 0.331 | 17.389 | 2157 |
| 1-Decanol | 0.297 | 15.578 | 1972 |
| Dimethyl sulfone | 0.293 | 9.035 | 1409 |
| Decyl octyl ether | 0.290 | 10.208 | 1499 |
| Propanoic acid, 2-methyl- | 0.284 | 11.048 | 1568 |
| Octadecanoic acid, butyl ester | 0.266 | 21.588 | 2645 |
| Propanoic acid, anhydride | 0.204 | 12.280 | 1671 |
| 1-Tridecene | 0.180 | 10.783 | 1547 |
| Phenol, 2,6-bis(1,1-dimethylethyl)-4-methyl-, methylcarbamate | 0.161 | 14.966 | 1913 |
| 2,4,7,9-Tetramethyl-5-decyn-4,7-diol | 0.158 | 16.755 | 2091 |
| cis-7,cis-11-Hexadecadien-1-yl acetate | 0.146 | 18.397 | 2266 |
| 2-Hexenal, (E)- | 0.138 | 6.469 | 1224 |
| 2,4,5-Trihydroxypyrimidine | 0.138 | 16.210 | 2035 |
| Formamide, N,N-dibutyl- | 0.137 | 13.562 | 1783 |
| Oleyl alcohol, trifluoroacetate | 0.135 | 25.591 | ** |
| Indolizine, 7-methyl- | 0.133 | 20.373 | 2495 |
| Phosphonofluoridic acid, methyl-, octyl ester | 0.130 | 12.218 | 1665 |
| Ethanol, 2-phenoxy- | 0.128 | 17.293 | 2147 |
| Cyclopentane, (1-methylethyl)- | 0.125 | 15.930 | 2006 |
| 1-Heptanol | 0.122 | 9.672 | 1459 |
| 2-Undecanone | 0.117 | 11.473 | 1601 |
| 2-Hexadecanone | 0.117 | 17.137 | 2131 |

| | | | |
|---|---|---|---|
| 1,12-Dodecanediol | 0.109 | 10.223 | 1500 |
| Benzene | 0.109 | 3.102 | 865 |
| Decanedioic acid, bis(2-ethylhexyl) ester | 0.106 | 29.129 | ** |
| 1,2-Diphenylcyclopropane | 0.105 | 20.238 | 2479 |
| Hexanedioic acid, bis(2-ethylhexyl) ester | 0.104 | 23.155 | 2832 |
| 2-Chloroethyl benzoate | 0.102 | 20.056 | 2457 |
| 2,4-Nonadienal, (E,E)- | 0.101 | 12.691 | 1705 |
| Tripropylene glycol monomethyl ether | 0.101 | 13.862 | 1810 |

**Supplementary Table S4: Triangle test with 54 panelists.**

| **Unalike Triangle Sample** | **Number of Correct Responses** |
|---|---|
| Pork Belly Fat | 20/24 |
| Cultured Pork Fat | 21/30 |
| Total | 41/54$^{\alpha}$ |

$^{\alpha}$ Critical number of correct responses required to demonstrate statistical significance ($p<0.05$) in a triangle test is 25/54 [40].

**Supplementary Table S5: Full triangle test summary**

Response of 1 indicates successful discrimination while 0 indicates unsuccessful discrimination between the samples. N/A indicates no answer was given.

| Panelist | Discrimination |
|---|---|
| 1 | 1 |
| 2 | 0 |
| 3 | 1 |
| 4 | 1 |
| 5 | 1 |
| 6 | 1 |
| 7 | 0 |
| 8 | 1 |
| 9 | 1 |
| 10 | 1 |
| 11 | 1 |
| 12 | 1 |
| 13 | 0 |
| 14 | 1 |
| 15 | 1 |
| 16 | 1 |
| 17 | 1 |
| 18 | 1 |
| 19 | 1 |
| 20 | 1 |
| 21 | 1 |
| 22 | 0 |
| 23 | 0 |
| 24 | 1 |
| 25 | 0 |
| 26 | 1 |
| 27 | 1 |
| 28 | 1 |
| 29 | 0 |
| 30 | 1 |
| 31 | 1 |
| 32 | 1 |
| 33 | 1 |
| 34 | 1 |
| 35 | 0 |
| 36 | 1 |
| 37 | 0 |
| 38 | 1 |

| | |
|---|---|
| 39 | 1 |
| 40 | 0 |
| 41 | 0 |
| 42 | 0 |
| 43 | 1 |
| 44 | 1 |
| 45 | 1 |
| 46 | 1 |
| 47 | 1 |
| 48 | N/A |
| 49 | 1 |
| 50 | 1 |
| 51 | 1 |
| 52 | 1 |
| 53 | 1 |
| 54 | 0 |
| 55 | 1 |

**Supplementary Table S6. Panelist demographics as percent of overall study**

| Age | | | Gender | | |
|---|---|---|---|---|---|
| 18 to 24 | 25 to 34 | 35 to 47 | Female | Male | Non-binary |
| 59.3% | 35.2% | 5.6% | 47.3% | 47.3% | 5.5% |

| Highest Level of Education | | | | | |
|---|---|---|---|---|---|
| High School | Some College | Technical School | Bachelor’s | Master’s | Doctorate |
| 7.3% | 32.7% | 0.0% | 45.5% | 7.3% | 5.5% |

| Annual Household Income | | | | | | | | |
|---|---|---|---|---|---|---|---|---|
| Under $20,000 | $20,000 - $39,999 | $40,000 - $59,999 | $60,000 - $79,999 | $80,000 - $99,999 | $100,000 - $119,999 | $120,000 - $139,999 | $140,000 - $159,999 | $160,000 and over |
| 13.5% | 11.5% | 26.9% | 5.8% | 5.8% | 7.7% | 1.9% | 3.8% | 23.1% |

| **Would you be willing to buy cultivated meat if it has the same price as conventional meat?** | **Would you be willing to pay 10% more for cultivated meat?*** | **Would you be willing to buy cultivated meat at a 10% discount?**** |
|---|---|---|
| Yes: 98.2% | Yes: 81.5% | Yes: 100.0% |
| * = out of participants who would buy cultivated meat at the same price as conventional meat.<br>** = out of participants who would **not** buy cultivated meat at the same price as conventional meat. | | |
